# Hydrogel confinement enables subcutaneous delivery of vesicant antibody-drug conjugates

**DOI:** 10.64898/2026.05.21.726796

**Authors:** Guillaume Jacquot, Capucine Hervy, Elies Belarouci, Adeline Gasser, Chawin Hoernel, Olga Traissard, María Gutiérrez-Blanco, Julien Draussin, Stéphane Erb, Susana Brun, Lyne Fellmann, Joris Mallard, Elyse Hucteau, Pierre Coliat, Stéphane Bernhard, Mark W Tibbitt, Jérôme Combet, Julien Graff, Maria Cristina Antal, Alain Carvalho, Laurent David, Céline Mirjolet, Sarah Cianferani, François Lux, Olivier Tillement, Sébastien Harlepp, Coralie Grange, Xavier Pivot, Alexandre Detappe

## Abstract

Antibody-drug conjugates (ADCs) deliver cytotoxic payloads to tumors with antibody selectivity, yet all approved ADCs are administered by intravenous (IV) infusion despite a strong patient and clinical preference for subcutaneous (SC) delivery. SC administration would reduce treatment burden, but many ADC payloads are vesicants that cause tissue necrosis upon local release, a liability amplified, not mitigated, by the dispersion-enhancing excipients used for SC antibody formulations. We developed an injectable diacetyl-L-tartaric anhydride-functionalized chitosan hydrogel (TACT) that addresses this conflict by confining ADCs within a protective SC depot. TACT is compatible with clinically approved ADC formulations without drug-product modification and provides drug-to-antibody ratio (DAR)-dependent release kinetics that support a quantitative relationship with in vivo absorption timing. In direct comparison, recombinant human hyaluronidase (rHuPH20) co-formulated with vesicant ADCs caused severe tissue necrosis, whereas TACT prevented macroscopic injury while preserving antitumor efficacy comparable to intravenous dosing. TUNEL staining of injection sites showed that TACT attenuated peri-depot apoptotic injury 3-fold relative to T-DM1 alone and 2-fold relative to rHuPH20 co-formulation. In non-human primates, SC TACT achieved 78% relative bioavailability for total trastuzumab, reduced peak circulating T-DM1 catabolite (free DM1) exposure 7.6-fold compared to IV administration and produced only transient, self-resolving cutaneous reactions. These results identify depot-mediated confinement as a viable alternative to excipient-mediated dispersion for SC delivery of vesicant ADCs, demonstrated here for trastuzumab-based conjugates across two approved ADC drug products (T-DM1 and T-DXd, with non-cleavable MCC and cleavable peptide linkers), with supporting validation in a custom cleavable monomethyl auristatin E (MMAE) series. Additional validation with enfortumab vedotin (EV), a Nectin-4-targeting MMAE ADC, supported the applicability of this strategy beyond trastuzumab-based conjugates.

## Introduction

The shift from intravenous (IV) to subcutaneous (SC) administration has reshaped the delivery of monoclonal antibodies (mAbs) in oncology.^1–3^ Fixed-dose SC formulations of trastuzumab, rituximab and trastuzumab-pertuzumab now permit administration in minutes rather than hours^4–6^, improving patient convenience and reducing treatment burden. For patients receiving chronic oncologic therapy, SC delivery reduces chair-time, enables home administration, and decreases infusion-related adverse events^7^. This transition was enabled by co-formulation with recombinant human hyaluronidase (rHuPH20) that transiently degrades the hyaluronan matrix of the SC space while dispersing large protein volumes through the interstitium^8,9^. For mAbs, this strategy has proven both safe and effective, facilitating widespread adoption of SC delivery. Recent high-concentration formulation strategies, including microparticle suspensions and excipient-stabilized concentrates, are extending SC administration to additional mAbs^10–12^. None of these strategies physically separates the payload from surrounding tissue, however, and none has been deployed for vesicant conjugates.

Antibody-drug conjugates (ADCs) represent a logical next class of candidates for SC administration^13–15^. However, vesicant and irritant ADCs present a distinct safety constraint that this approach does not address. Their cytotoxic payloads, including maytansinoids, auristatins, and camptothecin derivatives, cause concentration-dependent tissue necrosis upon local extravasation^16^, an intrinsic liability that dispersion-enhancing excipients would amplify rather than mitigate. To date, more than 14 ADCs have received regulatory approval, and all are administered by IV infusion. The prevailing strategy for SC ADC delivery has been to apply the same dispersion strategy that succeeded for conventional mAbs. Although non-vesicant/non-irritant SC ADC formulations co-administered with rHuPH20 are entering clinical evaluation (NCT07015697 for trastuzumab deruxtecan), no strategy has yet resolved the tension between dispersion-mediated delivery and vesicant payloads^17^.

We reasoned that safe SC delivery of vesicant ADCs could be achieved through the opposite strategy to dispersion, by confining the conjugate within a localized depot that shields surrounding tissue from payload exposure while releasing intact ADC into the systemic circulation at a controlled rate. Such a depot must be injectable through standard needles, gel in situ, retain the ADC without compromising structural integrity, and achieve bioavailability sufficient for therapeutic efficacy.

Here we report TACT (diacetyl-L-tartaric anhydride-functionalized chitosan hydrogel technology), an injectable hydrogel that meets these requirements. Using two clinically approved trastuzumab-based ADCs (T-DM1, DAR ∼3.5, non-cleavable MCC-DM1 linker; T-DXd, DAR ∼8, cleavable peptide-DXd linker) alongside a custom trastuzumab-MMAE conjugate series (DAR 2-8) and unconjugated trastuzumab as reference, we show that TACT confinement prevents tissue necrosis, preserves antitumor efficacy, and defines the key parameters that govern ADC retention and diffusion beyond the SC injection site. We further evaluated enfortumab vedotin (EV, DAR □3.8, cleavable mc-vc-PABC-MMAE linker), a clinically approved Nectin-4-targeting ADC, as an orthogonal non-HER2 validation.

## Results and discussion

### TACT: a cargo-responsive injectable hydrogel

We synthesized TACT by functionalization of chitosan with diacetyl-L-tartaric anhydride in a water/propanediol mixture at room temperature (Fig. 1A and Supplementary Fig. 1). Tartrate functionalization introduces carboxylate groups (pKa ∼3-4), deprotonated at physiological pH along the polysaccharide backbone. Unlike unmodified chitosan, which precipitates at physiological pH, TACT forms a stable hydrogel^**18,19**^ through physical ionic crosslinks between residual protonated amines and tartrate carboxylates. The result is a single-component system that gels in situ without chemical crosslinkers, avoiding potentially harmful by-products^**20–22**^.

**Figure 1.**
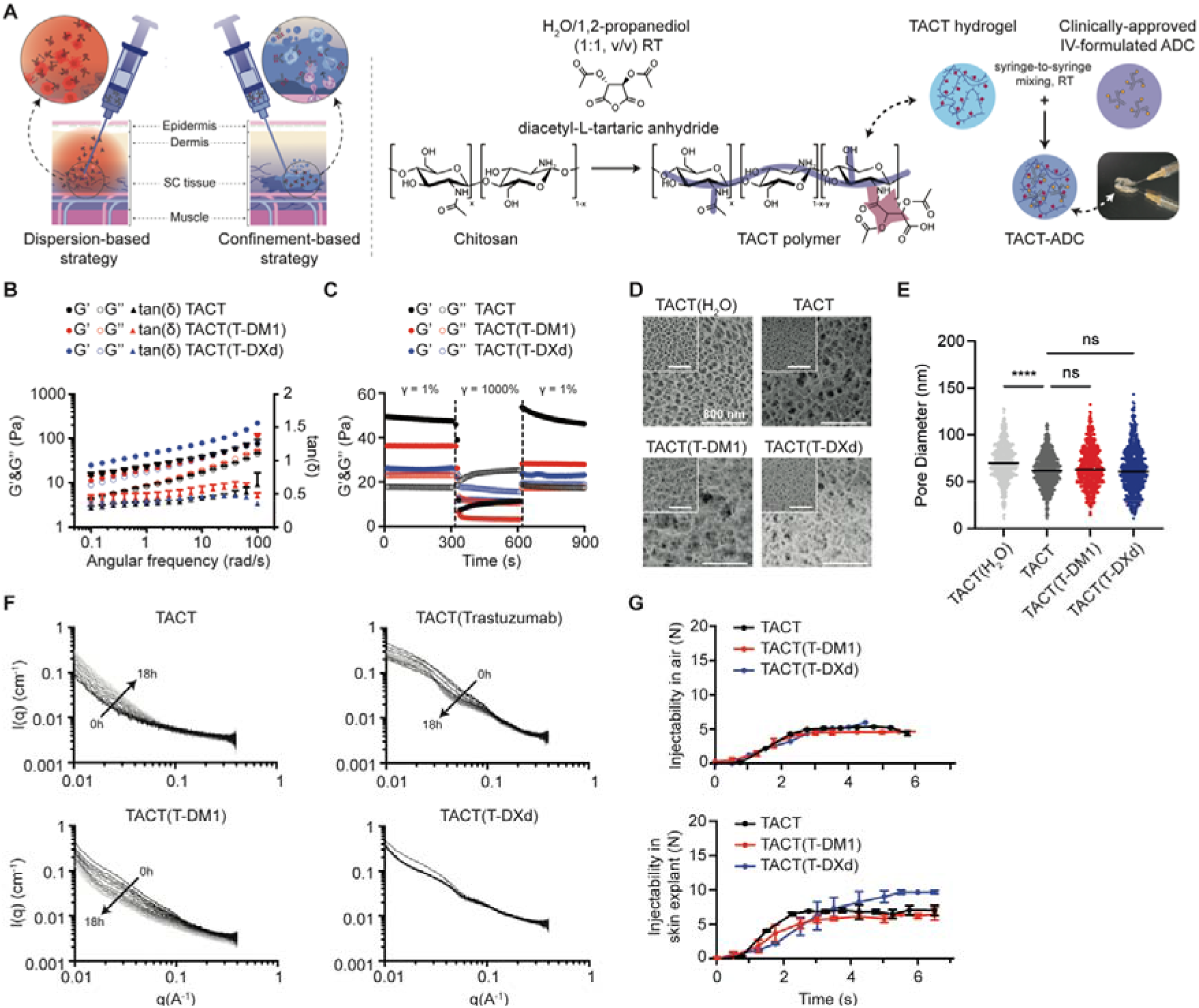
TACT, a cargo-responsive injectable hydrogel. **A**. Left: confinement strategy enabling slower drug-release kinetics and subcutaneous (SC) administration of vesicant drugs. Right: synthesis of TACT by functionalization of diacetyl-L-tartaric anhydride onto chitosan in a water/propanediol mixture at room temperature, and schematic of the syringe-to-syringe mixing process with clinically approved ADC formulations. **B**. Oscillatory rheology (frequency sweep) of TACT, TACT(T-DM1) and TACT(T-DXd) showing storage (G’, filled circle) and loss (G’’, open circle) moduli and tan delta (ratio G”/G’, triangle). **C**. Self-healing behavior assessed by cyclic strain, with recovery of G′ after repeated high-strain disruption. **D**. Representative cryo-SEM micrographs of TACT in water, physiological buffer (PBS 1x), and loaded with T-DM1 or T-DXd in PBS. Scale bar, 200 nm. **E**. Pore diameter distributions quantified from cryo-SEM (n = 1,761-3,112 measurements per condition). Horizontal lines indicate medians. P values from two-sided Mann-Whitney U tests with Bonferroni correction for six pairwise comparisons. **F**. Time-resolved SAXS profiles (I(q) versus q) for TACT alone, TACT(trastuzumab), TACT(T-DM1) and TACT(T-DXd) over 0-18 h. Greyscale gradient from black (t = 0) to light grey (t = 18 h). **G**. Extrusion force profiles at 1 mL/min for TACT, TACT(T-DM1) and TACT(T-DXd) through 25-gauge needles into air (upper part) and into full-thickness human skin explants (lower part). Data are mean ± SEM, **** indicates p-value <0.0001 using two-way ANOVA test.

TACT solutions (2.5-5% w/w) form optically transparent hydrogels (Fig. 1A). The storage moduli (G’; ω = 10 rad s^-1^) of unloaded TACT hydrogels increase with polymer concentration from G’ = 40 Pa for 2.5% w/w formulations to ∼150 Pa for 5% w/w formulations and exceed loss moduli (G”) across the frequency range tested (Fig. 1B, Supplementary Fig. 2 and Supplementary Table 1). Rheological properties were confirmed after co-formulation with approved ADC drug products: T-DM1 (ado-trastuzumab emtansine) and T-DXd (trastuzumab deruxtecan). Approved drug products were mixed with TACT by syringe-to-syringe transfer for 2 min without excipient modification. ADC loading up to 75 mg mL^-1^ in 2.5% w/w polymer formulations preserved gel-like behavior across the frequency range tested (Fig. 1B, Supplementary Fig. 2 and Supplementary Table 1). TACT gels exhibit robust shear-thinning and rapid elastic recovery (self-healing) with storage modulus recovering >70% of its initial value within seconds at low strain (γ = 1%) following high strain (γ = 1000%) disruption, indicative of the capacity of TACT hydrogels to form depots following injection (Fig. 1C). The shear-thinning and elastic recovery behavior is preserved in ADC-loaded TACT formulations.

Cryogenic scanning electron microscopy (cryo-SEM) reveals a porous microstructure whose dimensions are sensitive to ionic environment. In water, the TACT network exhibits a median pore diameter of 69 nm (interquartile range (IQR) 36-101; n = 2,704 measurements). Equilibration in physiological buffer (PBS 1x) reduces the median to 56 nm (IQR 29-90; n = 3,112), a contraction of 19% (P < 10^−15^, Mann-Whitney U test; Fig. 1D,E and Supplementary Table 2). Loading with T-DM1 or T-DXd does not alter the median pore diameter (P > 0.05 after Bonferroni correction). However, because an intact IgG (□5 nm hydrodynamic radius) is an order of magnitude smaller than the median pore, size exclusion alone cannot account for the observed cargo confinement. Time-resolved small-angle X-ray scattering (SAXS) further indicates that local network length scales can approach the IgG hydrodynamic radius (ξ = 5.7 nm for TACT(trastuzumab)), suggesting an additional contribution of steric hindrance and noncovalent gel-cargo interactions to early depot residence. Detailed structural characterization is provided in Supplementary Information.

Time-resolved SAXS (q = 0.005-0.40 Å^-1^) revealed that the TACT network reorganizes over hours in a cargo-dependent manner (Fig. 1F, Supplementary Fig. 3). Unloaded TACT densifies progressively, whereas in the presence of cargo three distinct trajectories emerged, with trastuzumab driving progressive network compaction, T-DM1 transiently disrupting the network before recovering to a compact state, and T-DXd reaching structural equilibrium within one hour without further reorganization. The resulting cargo-dependent structural hierarchy at 16 h (trastuzumab > T-DXd > T-DM1) does not follow DAR alone, indicating that payload chemistry and linker architecture modulate gel-conjugate interactions in addition to drug loading. Together, these observations define an active gel-cargo co-organization distinct from passive encapsulation, in which the antibody scaffold provides electrostatic contact sites for the chitosan-tartrate chains while attached payloads contribute hydrophobic associative patches (DM1 being more hydrophobic than DXd) whose nature and number jointly determine whether the network is disrupted, stabilized or compacted (complete time-resolved structural parameters and mechanism analysis are provided in the SI). Notably, scattering curves of cargo-loaded gels display two characteristic length scales, an inter-particle correlation (Θ) at low q and a chain-chain correlation length ξ at higher q, together with cargo-specific time evolution (Supplementary Fig. 3). This signature is consistent with a star-like local architecture in which the antibody acts as a multivalent core engaging chitosan-tartrate chains as arms, with the transient evolution of these length scales reflecting the progressive equilibration of antibody-polymer association. These structural measurements support cargo-dependent gel-cargo interactions but do not, on their own, predict release kinetics.

Having characterized cargo-dependent hydrogel structure, we next assessed whether TACT could accommodate approved ADC drug products spanning two clinically dominant linker chemistries: T-DM1 (Kadcyla, Roche; DAR □3.5, non-cleavable MCC-DM1 linker)^23,24^ and T-DXd (Enhertu, Daiichi Sankyo; DAR □8, cleavable peptide-DXd linker)^25,26^. All TACT(ADC) preparations remain injectable through 25-gauge needles, with extrusion forces below 10 N (Fig. 1G), well below the ∼20 N threshold commonly associated with successful manual injection.

### ADC integrity, controlled release, pharmacokinetics, and therapeutic efficacy

We first assessed whether gel residence alters ADC integrity. Size-exclusion chromatography coupled to native mass spectrometry (SEC-nMS) profiles, at both the intact and subunit (middle-up) levels, were comparable between released samples from gel and fresh controls (Fig. 2A-D, Supplementary Fig. 4 and Supplementary Table 3-8), confirming that TACT does not alter the drug product.

To determine whether tartrate functionalization is essential for cargo retention, we compared antibody release from TACT and unmodified chitosan in simulated SC fluid at room temperature.

Beyond polymer chemistry, release kinetics are governed by drug loading. To isolate the contribution of drug loading independent of linker chemistry and clinical excipients, we synthesized a series of trastuzumab-MMAE conjugates with DAR values of 0, 2, 3.5 and 8 using identical mc-vc-PABC-MMAE linker architecture, a cleavable linker chemistry validated in several approved ADCs including brentuximab vedotin and polatuzumab vedotin^27–29^ (Supplementary Fig. 5-6 and Supplementary Table 9). The released amount decreases monotonically with increasing DAR (cumulative release at 6 h: 19.4 ± 2.6% for DAR 0, 14.6 ± 2.3% for DAR 2, 12.6 ± 1.0% for DAR 3.5 and 11.1 ± 0.8% for DAR 8; Fig. 2E), establishing DAR as an intrinsic determinant of release independent of linker chemistry and clinical excipients. We next tested whether this DAR-release relationship was preserved in approved ADC drug products. Unconjugated trastuzumab (DAR 0) releases most rapidly from TACT, followed by T-DM1 (DAR ∼3.5), with T-DXd (DAR ∼8) releasing most slowly, with a cumulative release at 1 h of 35.1 ± 1.8%, 27.5 ± 1.3%, and 21.5 ± 0.3%, respectively. A similar trend is observed at 6 h with cumulative release of 63.6 ± 3.1% for trastuzumab, 65.5 ± 6.2% for T-DM1 and 50.9 ± 0.9% for T-DXd (Fig. 2F, Supplementary Fig 7). However, release from these marketed drug products is faster than from excipient-free conjugates at matched DAR, indicating that clinical formulation components modulate depot-cargo interactions in addition to ADC architecture. This effect is already evident at DAR 0, where clinical trastuzumab (Trazimera, Pfizer) releases 3.3-fold more at 6 h than the matched excipient-free antibody (63.6 versus 19.4%). Together, these data establish DAR as the dominant intrinsic determinant of depot retention when formulation is held constant, while identifying drug-product excipients as a powerful extrinsic modulator of release kinetics. Increasing buffer strength also progressively slowed T-DM1 release (Supplementary Fig. 8), indicating that ionic environment also modulates gel-cargo and gel-gel interactions in addition to DAR and payload chemistry. Importantly, this release hierarchy does not directly mirror the SAXS-derived structural hierarchy, indicating that DAR, payload chemistry and drug-product excipients dominate release behavior beyond global network architecture.

**Figure 2.**
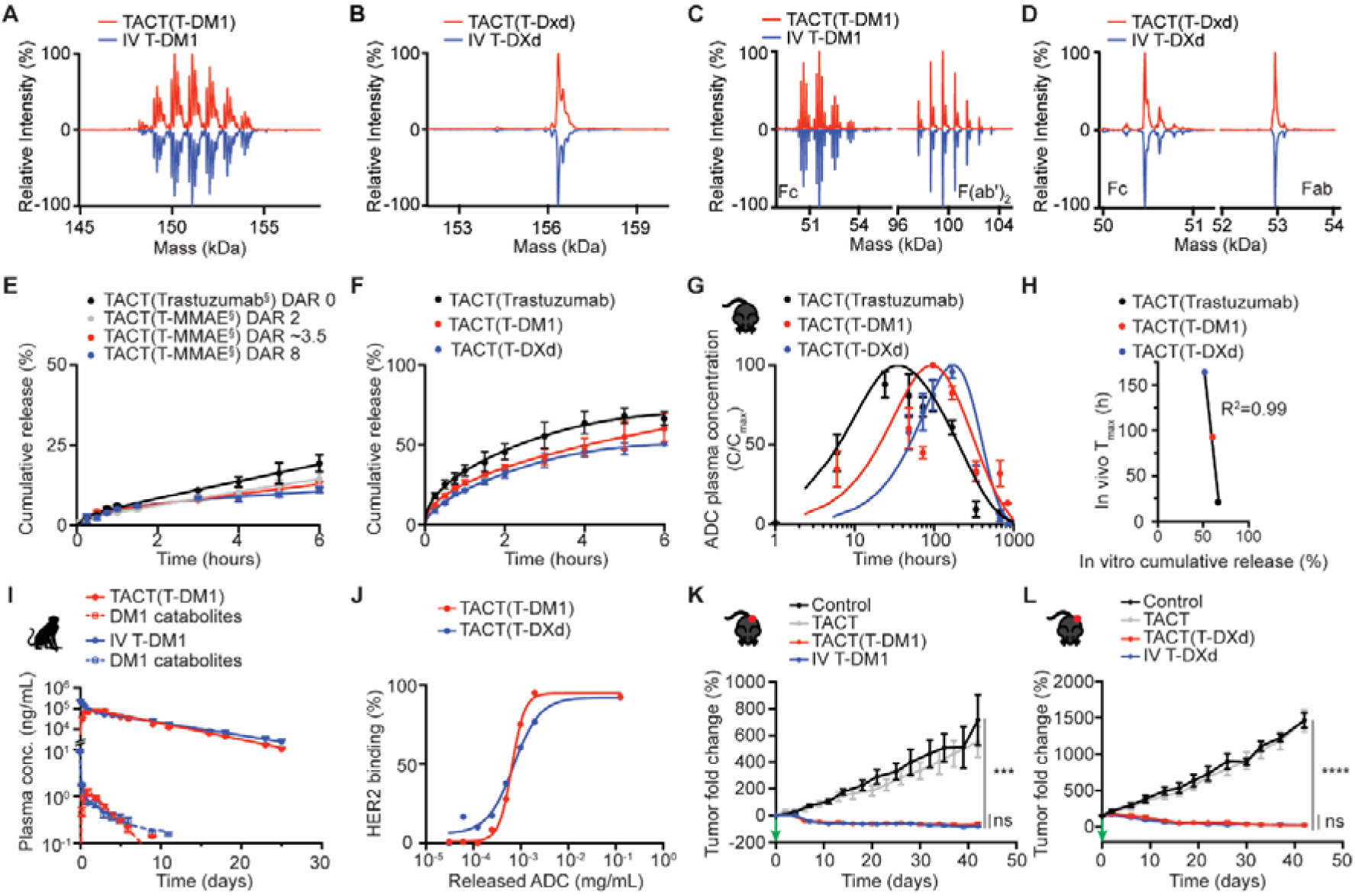
ADC integrity, controlled release, pharmacokinetics and therapeutic efficacy. **A-D**. ADC physicochemical integrity after gel residence assessed by intact SEC-nMS for T-DM1 (A) and T-DXd (B) and by middle-up SEC-nMS for T-DM1 (C) and T-DXd (D), TACT-released ADC (red), and reference IV formulation (blue). **E**. Cumulative in vitro release from TACT for excipient-free trastuzumab-MMAE conjugates (DAR 0, 2, 3.5, 8) in PBS. ^§^ Excipient-free. **F**. Cumulative release of approved drug products: trastuzumab (Trazimera), T-DM1 (Kadcyla) and T-DXd (Enhertu). **G**. Normalized (C/Cmax) plasma concentration-time profiles after SC TACT delivery of trastuzumab, T-DM1 and T-DXd, showing DAR-dependent delayed Tmax. **H**. In vitro-in vivo correlation (IVIVC): cumulative in vitro release at 6 h versus in vivo Tmax. R^2^ from linear regression. **I**. NHP pharmacokinetics in cynomolgus macaques (n = 3 per group). Plasma concentrations of intact T-DM1 (solid lines) and free DM1 catabolite (dashed lines) after IV (blue) or SC TACT (red) administration. **J**. HER2 binding of ADC recovered after TACT gel residence versus fresh controls, assessed on BT-474 cells. **K**. Tumor growth in BT-474 xenograft-bearing mice treated with SC TACT(T-DM1) or IV T-DM1 (10 mg kg^-1^; n = 8 per group). ns, not significant by two-tailed Mann-Whitney U test. **L**. Tumor growth in BT-474 xenograft-bearing mice treated with a single dose of SC TACT(T-DXd) or IV T-DXd (10 mg kg^-1^; n = 8 per group). Green arrow indicates treatment day. Data are mean ± SEM, ***P < 0.001, ****P < 0.0001 versus control; ns, not significant between SC and IV groups.

Biexponential modeling of the MMAE DAR series confirmed two-phase release kinetics (Supplementary Fig. 9 and Supplementary Table 10), with a fast initial burst and a slower sustained release phase governing the majority of payload delivery. Complementary Korsmeyer-Peppas analysis^30^ of the early release window (Q ≤ 60%) provided mechanistic insight into the transport regime. The release exponent n decreased with increasing DAR (0.68 ± 0.04 for DAR 0, 0.70 ± 0.03 for DAR 2, 0.50 ± 0.03 for DAR 3.5, 0.48 ± 0.04 for DAR 8), revealing a transition from anomalous transport (n ≈ 0.7, coupled diffusion and polymer relaxation) at low DAR to essentially Fickian diffusion (n ≈ 0.5) at high DAR. This shift indicates that above DAR ∼2, payload hydrophobicity strengthens gel-conjugate interactions sufficiently to make diffusion through the network the rate-limiting step. The shift in best-fit model observed at DAR 8, from a biphasic diffusion-relaxation regime to a diffusion-from-sphere model with a finite releasable fraction, further indicates that the physicochemical properties of the conjugate at high drug loading alter the predominant transport mechanism, resulting in both a reduced release rate constant (k_BL_ = 0.0045 h^-1^) and an incomplete release plateau under the experimental conditions tested. Collectively, these data establish DAR as a primary determinant of MMAE release kinetics from the TACT hydrogel depot, with a progressive transition from relaxation-coupled diffusion to network diffusion-limited release as DAR increases.

Following SC delivery in TACT, systemic concentrations in healthy C57BL/6 mice increase gradually. When concentration-time curves were normalized to C_max_, higher DAR was associated with delayed T_max_ across cargos (Fig. 2G and Supplementary Table 11). Noncompartmental analysis of mean concentration-time profiles reveals a DAR-dependent pharmacokinetic profile hierarchy. For T-DM1 (DAR □3.5; 0.35 mg IV and SC), SC TACT delivery yields a relative bioavailability of 61%, with C_max_ of 254 versus 350 mg L^-1^ for IV dosing (1.4-fold reduction), T_max_ of 96 h and a terminal half-life of 333 h. For T-DXd (DAR □8; 0.20 mg IV and SC), bioavailability decreases to 36% and C_max_ is lower by 2.7-fold (74 versus 200 mg L^-1^), with T_max_ of 168 h, reflecting prolonged depot retention of the highest-DAR conjugate. The lower bioavailability and shorter apparent terminal half-life of T-DXd after SC TACT administration indicate stronger depot retention of the high-DAR conjugate and a smaller systemically available released fraction. The in vitro release hierarchy was preserved in vivo, with trastuzumab releasing fastest, followed by T-DM1 and then T-DXd. DAR correlated positively with T_max_ (n = 3, P = 0.046), and cumulative in vitro release at 6 h correlated inversely with in vivo T_max_ (Spearman rs = -1.0, n = 3, Fig. 2H), supporting a quantitative in vitro-in vivo relationship for TACT-mediated depot release.

We confirmed the translational relevance of depot-mediated release in non-human primates (NHPs). Cynomolgus macaques were randomized into three groups (n = 3 per group): IV T-DM1 (10 mg kg^-1^), SC TACT alone (650 µL) and SC TACT(T-DM1) (650 µL, ∼10 mg kg^-1^). Pharmacokinetic analysis of T-DM1 revealed a depot-release profile characterized by delayed T_max_ (64 ± 8 h) and lower C_max_ (82.9 ± 13.6 versus 232.5 ± 19.4 µg mL^-1^ for IV dosing), with preserved total systemic exposure (AUC_0−∞_; relative bioavailability 78.1 ± 20%; Fig. 2G and Supplementary Fig. 10 and Supplementary Table 12). Plasma concentrations of free DM1, quantified by LC-MS/MS, reveal marked attenuation of peak free payload exposure after SC TACT delivery. IV T-DM1 produced a DM1 C_max_ of 9.1 ng mL^-1^, which SC TACT reduced 7.6-fold to 1.2 ng mL^-1^ (Fig. 2I). Total DM1 catabolite exposure is preserved (AUC_0−∞_ 4.1 versus 5.4 ng day mL^-1^; 76% of IV exposure), reflecting attenuated early systemic catabolite burst rather than reduced overall deconjugation. Across the three analytes measured, the depot preferentially attenuates peak exposure in proportion to analyte toxicity. C_max_ reduction is 3.0-fold for total trastuzumab, 2.8-fold for intact T-DM1, and 7.6-fold for free DM1. This selective attenuation of the catabolite most directly associated with dose-limiting toxicity provides preclinical evidence for a favorable translational safety profile of SC TACT delivery. For several ADCs, peak exposure is associated with dose-limiting toxicities, including hepatotoxicity and thrombocytopenia for T-DM1^31,32^ and interstitial lung disease and cardiotoxicity for T-DXd^33^. The 2.8-fold reduction in T-DM1 C_max_ and 7.6-fold reduction in free DM1 C_max_, together with preserved total systemic exposure, provide preclinical rationale for a pharmacokinetic profile that may decouple efficacy from peak-driven toxicity.

A major question is whether depot-mediated release preserves therapeutic efficacy despite the altered pharmacokinetic profile. In vitro, we first assessed HER2 binding using HER2-positive BT-474 cells (Fig. 2J), in agreement with the physicochemical integrity data and confirming that gel-cargo interactions do not alter antigen recognition. In mice bearing BT-474 xenografts, SC TACT(T-DM1) achieves tumor growth inhibition indistinguishable from IV T-DM1 in both single-dose and multi-dose (Q7Dx4) regimens (P = 0.92, two-tailed Mann-Whitney U test; Fig. 2K, Supplementary Fig. 11). A single SC injection of TACT(T-DXd) (10 mg kg^-1^) induced tumor regression comparable to IV T-DXd in BT-474 xenografts (n = 8 per group, Fig. 2L), extending the platform across non-cleavable MCC and cleavable peptide linker conjugates. We further extended platform versatility to a non-HER2 backbone with enfortumab vedotin (EV), a Nectin-4-targeting MMAE ADC^27,28^. TACT(EV) preserved hydrogel rheology and shear-thinning behavior at 2.5% w/w polymer with 50 mg mL^-1^ EV, released EV progressively in simulated SC fluid, prevented the macroscopic tissue injury observed with EV alone, and achieved antitumor efficacy comparable to IV EV in HT-1376 xenografts after a single 10 mg kg^-1^ dose (n = 8 per group, P < 0.0001 versus control, not significant between SC TACT(EV) and IV EV; Supplementary Fig. 18).

### Confinement prevents dispersion-driven tissue injury and supports non-human primate translation

To test whether confinement and dispersion produce distinct safety outcomes for vesicant payloads, we compared four SC T-DM1 administration strategies in C57BL/6 mice: (i) TACT(T-DM1) hydrogel, (ii) TACT alone (vehicle control), (iii) rHuPH20 co-formulation at 1,000 U, and (iv) T-DM1 in its approved IV formulation excipients without additional excipient. The rHuPH20-T-DM1 preparation remained visually clear before injection, ruling out gross aggregation as a confounder. As expected for a vesicant payload, T-DM1 alone produces moderate erythema and induration peaking at day 3, consistent with clinical extravasation guidance^***34***^. By contrast, rHuPH20 co-formulation markedly exacerbated local injury. Extensive necrosis appears within 72 h and progresses to full-thickness ulceration by day 5 (Fig. 3A). Histopathology reveals widespread tissue degeneration, inflammatory infiltration, and vascular damage extending well beyond the injection site (Fig. 3B). The spatial pattern of injury, radiating outward from the injection locus, is compatible with enzymatic degradation of the hyaluronan barrier promoting ADC dispersion through tissue planes and thereby exposing a larger volume of SC tissue to cytotoxic payload. By contrast, TACT(T-DM1) produces no overt macroscopic necrosis. Anti-human Fc immunohistochemistry confirms that the conjugate remains confined within the gel depot rather than dispersing through surrounding tissue (Fig. 3C). In rHuPH20-treated sites, tissue disruption limited interpretation of focal depot-associated staining, but the remaining anti-trastuzumab signal appeared broader and less structured than in TACT(T-DM1) sections. Body weights are stable across all groups (Fig. 3E), indicating that injury was local rather than systemic.

**Figure 3.**
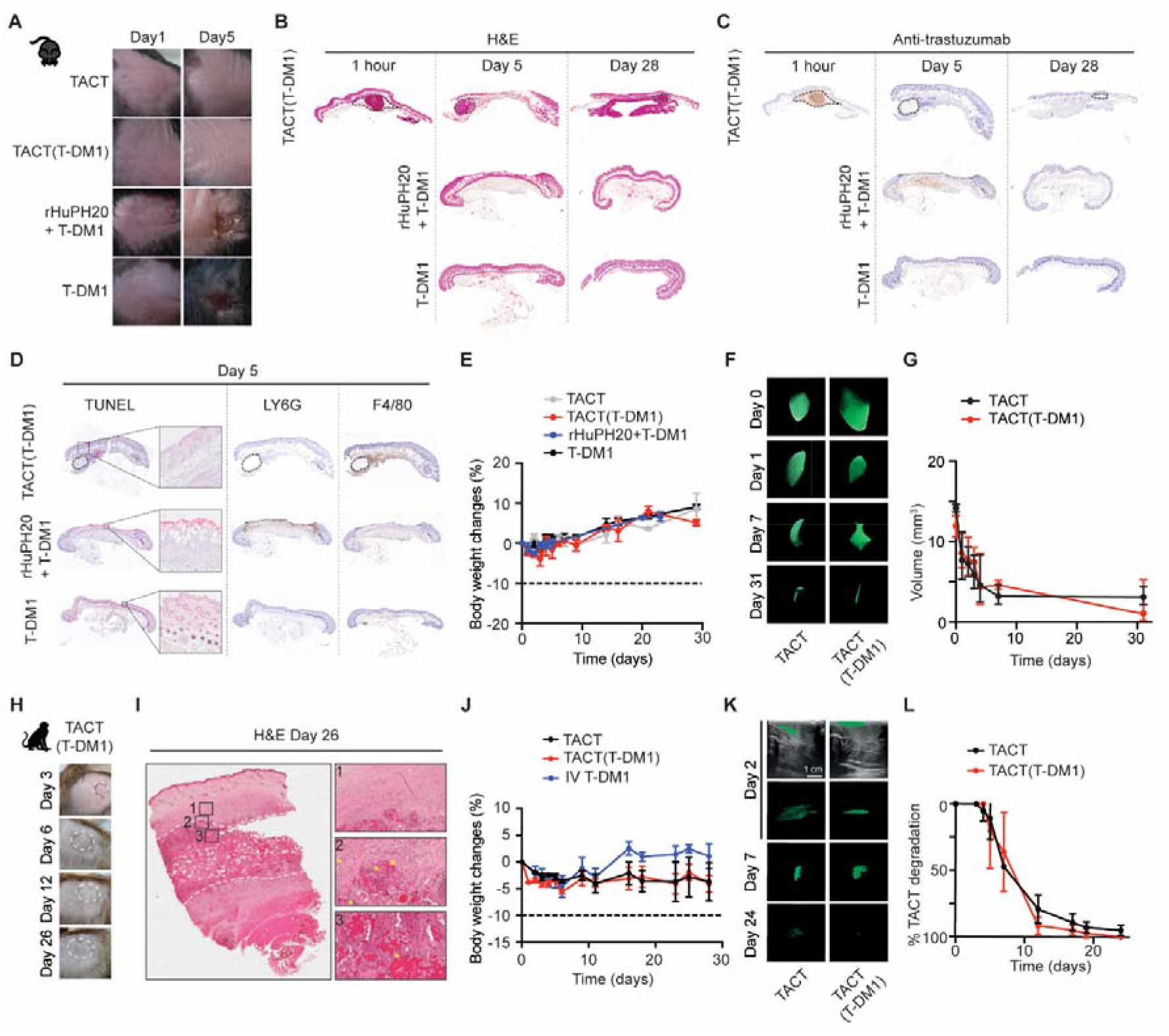
Confinement prevents dispersion-driven tissue injury and supports NHP translation. **A**. Macroscopic photographs of SC injection sites in C57BL/6 mice at days 1 and 5 for TACT alone, TACT(T-DM1), T-DM1 alone, and rHuPH20 + T-DM1, at a similar injected dose of 30 mg kg^-1^ TDM1. **B**. Hematoxylin and eosin (H&E) staining of SC injection sites in C57BL/6 mice at 1 hour, day 5 and day 28 for TACT(T-DM1), rHuPH20 + T-DM1, and T-DM1 alone. **C**. Anti-trastuzumab immunohistochemistry of SC injection sites in C57BL/6 mice at 1 hour, day 5 and day 28 for TACT(T-DM1), rHuPH20 + T-DM1, and T-DM1 alone. **D**. TUNEL, F4/80+ and Ly6G+ immunostaining of SC injection sites at day 5 for TACT(T-DM1), rHuPH20 + T-DM1, and T-DM1 alone. **E**. Body weight changes (%) over 30 days in the local tolerance study (n = 5 per group). Dashed line indicates 10% body weight loss limit. **F**. Representative MRI three-dimensional segmentation of SC depots (green) for TACT alone and TACT(T-DM1) at days 0, 1, 7 and 31. Scale bar, 2 cm. **G**. Depot volume quantified by MRI segmentation over 31 days (n = 3 per group). **H**. Macroscopic photographs of NHP injection sites at days 3, 6, 12 and 26. **I**. End-of-study (day 26) histopathology (H&E) of NHP injection sites with numbered insets showing depot-tissue interface regions: (1) collagen shell, (2) collagen shell-TACT interface, (3) TACT hydrogel. **J**. Body weight changes (%) of NHPs over the treatment period for TACT alone, TACT(T-DM1) and IV T-DM1 (n = 3 per group). Dashed line indicates 10% body weight loss limit. **K**. Representative ultrasonography images of SC depots in cynomolgus macaques at days 2, 7 and 24. Scale bar, 1 cm. **L**. TACT degradation kinetics quantified by 3D segmentation of serial ultrasonography data (n = 3 per group). Data are mean ± SEM

To resolve how delivery mode shapes tissue-level ADC exposure following SC administration, we performed anti-trastuzumab immunohistochemistry on injection sites over a time course spanning the acute (1 h, day 1), intermediate (day 4, day 7) and resolved (day 28) phases of depot evolution for TACT(T-DM1) (Supplementary Fig. 19). TACT(T-DM1) retained anti-trastuzumab signal within the depot at early time points, with minimal extension into surrounding tissue at 1 h and day 1, followed by progressive redistribution from the depot periphery at day 4, and day 7 (Fig. 3C), consistent with controlled release into the adjacent interstitium. T-DM1 alone produced an intermediate distribution without a persistent focal depot, whereas rHuPH20 co-formulation produced a broader and less structured tissue distribution, consistent with dispersion-mediated spread through the subcutaneous space (Fig. 3C). By day 28, anti-trastuzumab staining in TACT(T-DM1) sections was faint and diffuse, consistent with clearance of released ADC concurrent with depot resorption (Fig. 3). The distinct exposure patterns observed by immunohistochemistry translated into quantitatively distinct cytotoxic activity at the tissue level. TUNEL staining at day 5, the comparative peak-injury timepoint for T-DM1 alone and rHuPH20 co-formulation, revealed that TACT(T-DM1) reduced peri-depot apoptotic signal 3-fold relative to T-DM1 alone (1.39% versus 4.19% of tissue area) and 2-fold relative to rHuPH20 co-formulation (1.39% versus 2.77%; Fig. 3D). Beyond amplitude, TACT(T-DM1) also altered the kinetics of local cytotoxicity. Apoptotic signal followed a delayed, multiphasic trajectory (0.14% at 1 h, 0.49% at day 1, 0.24% at day 5) before resolving to near-baseline by day 28 (0.15%) (Supplementary Fig. 20). Together, these data indicate that hydrogel confinement does not abolish local pharmacological activity but attenuates its peak amplitude, delays its onset, and accelerates its resolution relative to dispersion-mediated delivery.

To characterize the host response to each delivery mode, we quantified F4/80+ macrophage and Ly6G+ neutrophil infiltration at the injection site (Fig. 3D). At day 5, TACT(T-DM1) reduced intradepot neutrophil density 35-fold relative to rHuPH20 co-formulation (4.3 versus 150.1 cells mm^-2^), with immune cells distributed across the surrounding tissue rather than accumulating at the depot margin as observed under rHuPH20. Within TACT(T-DM1) depots, neutrophil density peaked transiently at day 1 and returned to baseline by day 5, while macrophage density increased from day 1 to day 5 before decreasing at day 28, a temporal sequence consistent with progressive biomaterial resorption (Supplementary Fig. 20). The acellular gel depot physically separates ADC from viable tissue during the early high-concentration window, whereas rHuPH20-mediated dispersion exposes the conjugate directly to SC tissue planes. These findings show that SC delivery strategies validated for benign antibodies cannot be safely transposed to vesicant ADCs.

Longitudinal magnetic resonance imaging (MRI) visualized depot behavior in vivo (Fig. 3F,G). Three-dimensional segmentation confirms well-defined SC depots at day 0 (approximately 15 mm^3^ for TACT alone and 10 mm^3^ for TACT(T-DM1)) that undergo biphasic resorption, with rapid volume reduction within the first 7 days (to ∼3-4 mm^3^), followed by slower clearance and near-complete resorption by day 31 (Fig. 3F,G). Both TACT alone and TACT(T-DM1) depots followed this superimposable biphasic clearance profile, indicating that ADC loading does not alter hydrogel resorption kinetics. Hematoxylin and eosin staining shows preserved tissue architecture surrounding the depot at all time points (Supplementary Fig. 14). The majority of ADC cargo exits the depot within 24-72 h, whereas the hydrogel matrix persists for approximately 25-31 days, a temporal asymmetry that provides physical containment during the critical early window of highest local ADC concentration, after which the gel resorbs as an inert, payload-free biomaterial.

Having established injectability in human skin explants (Fig. 1G), we next evaluated SC TACT(T-DM1) in NHPs. Cynomolgus macaques (n = 3 per group) received either IV T-DM1 (10 mg kg^-1^), SC TACT alone (650 µL) or SC TACT(T-DM1) (650 µL of 50 mg mL^-1^ T-DM1 in 2.5% w/w TACT; ∼10 mg kg^-1^). Injection was completed in approximately one minute through 25-gauge needles. Given the severe tissue necrosis observed in mice, SC administration of T-DM1 alone or with rHuPH20 was deemed ethically unacceptable for NHP evaluation. Animals in the TACT(T-DM1) group developed transient cutaneous reactions at injection sites (erythema and edema beginning at day 2, with desquamation peaking around days 5-7) that were managed with topical supportive care and resolved completely by day 21 (Fig. 3H). No such reactions occurred in the TACT alone group, indicating that the local response was attributable to the ADC payload rather than the hydrogel vehicle, consistent with the known vesicant profile of T-DM1. Body weights, vital signs and clinical biochemistry remained within physiological ranges throughout the 29-day observation period (Fig. 3J and Supplementary Fig. 15 and 16). C-reactive protein levels remained low (≤1.4 mg L^-1^ at endpoint), confirming no systemic inflammatory response. End-of-study histopathology at day 29 shows ongoing hydrogel remodeling with immune-cell infiltration observed by H&E staining but no residual necrosis, fibrosis or cytosteatonecrosis at any injection site (Fig. 3H,I). This safety profile, characterized by transient cutaneous reactions without deep tissue injury and distinct from dispersion-based delivery, supports the preclinical feasibility of SC delivery for vesicant ADCs.

Serial ultrasonography of NHP injection sites tracked depot persistence over 24 days. Depots were clearly visible at day 2, progressively reduced in volume by day 7, and nearly undetectable by day 24 (Fig. 3K). Quantitative analysis reveals sigmoidal clearance kinetics, with progressive depot resorption between days 2 and 24 consistent with the slower turnover observed by H&E histology at day 29 (Fig. 3I, Fig. 3L). Degradation profiles of TACT alone and TACT(T-DM1) are superimposable, confirming that the initial presence of the ADC does not alter hydrogel clearance.

Although chitosan has an established biocompatibility record, the tartrate modification introduces a moiety whose immunogenic potential has not been formally assessed. Chitosan hydrogels are expected to undergo in vivo degradation by lysozyme and tissue glycosidases into low-molecular-weight chitooligosaccharides amenable to renal clearance^35^ and repeated SC dosing (Q7D×4) in mice produces no cumulative local toxicity or loss of efficacy (Supplementary Fig. 17). Together with the absence of systemic inflammatory signal in non-human primates, these data support the local and systemic tolerability of repeat TACT administration, although extended immunogenicity assessment in humans will be required.

## Conclusion

The transition from IV to SC delivery is already reshaping antibody therapeutics, and ADCs represent a logical next candidate for this shift. In this work, we report two advances relevant to SC ADC delivery. First, we show that dispersion of vesicant conjugates throughout tissue compromises local safety. Co-formulation with rHuPH20 results in extensive necrosis from peak tissue ADC exposure during the early dispersion phase. At the cellular level, peri-depot apoptotic injury was 3-fold lower with TACT than with T-DM1 alone and 2-fold lower than with rHuPH20 co-formulation at the peak timepoint, with apoptotic signal resolving to baseline by day 28.

TACT mitigates this local toxicity by confining vesicant ADCs within an acellular depot that shields surrounding tissue during the critical early window. This confinement principle enables safe SC delivery without redesign or re-engineering of the ADC itself. Approved IV drug products are converted to SC depots through simple syringe mixing, preserving the manufacturing, supply chain and regulatory pathways already established for the parent products. Confinement, rather than dispersion, is the enabling principle for SC delivery of vesicant ADCs, and TACT broadens access to a safer and more convenient route of administration that benefits patients, oncology nurses and healthcare systems alike.

Second, we show that ADC depot residence and release are governed jointly by DAR, payload chemistry, drug-product excipients and cargo-dependent gel-cargo interactions. The observation that clinical drug-product excipients substantially modulate depot release identifies formulation composition as a further design axis for confinement-based delivery. These findings distinguish the SC delivery requirements of unconjugated antibodies from those of vesicant ADCs, a distinction not addressed by current SC strategies. Consequently, the transition from IV to SC ADC delivery will require payload-aware formulation design rather than a simple adaptation of antibody-based SC strategies. Across ADCs in which peak systemic or local exposure contributes to dose-limiting toxicity, including T-DM1 and the broader class of MMAE- and DXd-based conjugates, confinement provides a practical route to convert IV drug products into SC depots without modifying the conjugate. Depot residence is governed by DAR, payload/linker chemistry and clinical formulation excipients, and can be prioritized through simple release, injectability and ADC-integrity assays. These findings identify irritant/vesicant ADCs, including T-DM1 and EV, as leading candidates for clinical translation of confinement-based SC ADC delivery. Beyond ADCs, the same confinement principle may be applicable to other narrow therapeutic-index biologics whose peak systemic or local exposure constrains dosing, opening a route toward home-administered, patient-friendly delivery for the next generation of injectable cancer therapeutics.

## Materials and Methods

All chemicals, polymers and reagents were obtained from commercial suppliers and used as received unless otherwise noted. Trastuzumab (Trazimera, Pfizer), trastuzumab emtansine (T-DM1, Kadcyla, Roche) and trastuzumab deruxtecan (T-DXd, Enhertu, Daiichi-Sankyo) were supplied as clinical lyophilized vials by the pharmacy department of the Institut de Cancérologie Strasbourg Europe and reconstituted according to manufacturers’ instructions.

### Synthesis of TACT polymer

TACT was synthesized by functionalization of (+)-diacetyl-L-tartaric anhydride onto raw chitosan (Heppe Medical Chitosan, degree of acetylation 2.6 ± 0.6%) in a water/propanediol mixture (1:1, v/v) at room temperature. The product was purified by tangential-flow filtration (50 kDa MWCO, 15 diavolumes) and freeze-dried. Functionalization was confirmed by ^1^H NMR (degree of functionalization 25 ± 1%) and FTIR. Full synthesis and characterization details are provided in Supplementary Information.

### Preparation of TACT hydrogels and loading of ADCs

Lyophilized TACT was dissolved at 10% (w/w) in water (pH 5.7), transferred to Luer-lock syringes, centrifuged to remove air bubbles and sterilized by autoclaving (121°C, 20 min). Clinically approved formulations were mixed with TACT by syringe-to-syringe transfer (200 reciprocations, 2 min) to obtain homogeneous hydrogels with final antibody concentrations of 50 mg mL^-1^ in 2.5% w/w TACT.

### In vitro release assays

TACT hydrogels loaded with antibody or ADC (100 µL) were injected into 1.5 mL tubes containing 1 mL of simulated SC fluid (pH 7.4, 25°C). At predefined time points (0.25-6 h), 2 µL of supernatant were withdrawn in triplicate and concentrations quantified by UV absorbance at 280 nm. Release kinetics were modeled using Korsmeyer-Peppas, Higuchi, first-order, Weibull and Hixson-Crowell equations with model selection by AIC.

### Small-angle X-ray scattering (SAXS)

SAXS data were collected on a Xeuss instrument with Cu Kα radiation (λ = 1.542 Å) at a sample-to-detector distance of 600 mm, covering q = 0.005-0.40 Å^-1^. Hydrogels were mounted in 1.008 mm-thick capillaries and measured every 30 min over 17 h. Extended kinetics (0-98 h) were collected on the DiffériX platform (Institut Charles Sadron). Data were analyzed using a custom Python pipeline fitting four complementary structural models to each I(q) spectrum: (i)□a Guinier analysis (q□=□0.005-0.030□Å^-1^, validity q.Rg□< □1.3) extracting the radius of gyration Rg; (ii) □an Ornstein-Zernike model (I(q) □= □I_0_/(1□+ □q^2^ξ^2^), q□=□0.010-0.065□Å^-1^) providing the correlation length ξ as a local mesh-size proxy; (iii)□a generalized Porod fit (I(q) □∼□q^-n^, q□=□0.060-0.180□Å^-1^) reporting local chain flexibility; and (iv)□a mass-fractal fit (q□=□0.020-0.100□Å^-1^) yielding the fractal dimension D_fractal_. The power-law exponent α reported in the Results equals −D_fractal_ from model (IV). All model fits required R^2^□>□0.80; fits failing this criterion were flagged and excluded from kinetic analyses.

### Mouse pharmacokinetic studies

All mouse experiments were approved by the relevant institutional animal care and use committee (APAFIS number #47391-2024020517379486, #38306-2022082410083076, #45506-2023102611073304). Female C57BL/6J or BALB/c mice (16-25 g) received SC injections of TACT(ADC) (2.5% w/w, 50 mg mL^-1^) in the dorsal flank using 30-gauge needles, or IV injections via the tail vein. Blood was collected at serial time points into EDTA-coated tubes. Plasma trastuzumab concentrations were quantified using a TR-FRET assay (HTRF Human IgG Detection kit, Cisbio/Revvity) or TR-FLISA assay (Poly-Dtech). Noncompartmental pharmacokinetic parameters were derived from mean concentration-time profiles.

### Tumor models and efficacy studies

BT-474 cells (2×10^7^) in 50% (v/v) Matrigel were implanted SC in female athymic nude mice (n = 8 per group). Treatment began at □160 mm^3^. For T-DM1: Q7Dx4 at 5 mg kg^-1^ comparing IV Kadcyla with SC TACT(T-DM1). For T-DXd: single dose at 10 mg kg^-1^. Tumor volumes (length × width^2^/2) and body weights were measured at least twice weekly.

### Local tolerance and histology

C57BL/6 mice received SC injections of T-DM1 (Kadcyla), TACT(T-DM1), T-DM1 + rHuPH20 (1,000 U) or TACT alone. Injection sites were inspected daily and photographed. Biopsies were collected at predetermined time points, fixed in 4% paraformaldehyde, embedded in paraffin and sectioned at 5 µm. H&E staining, anti-human Fc and anti-trastuzumab immunohistochemistry, TUNEL labeling, and immunostaining for F4/80 (macrophages) and Ly6G (neutrophils) were performed on serial sections. Slides were examined by a blinded pathologist. Whole-slide images were quantified using automated thresholding to determine percent TUNEL+ tissue area, and immune-cell densities (F4/80+ and Ly6G+ cells per mm^2^) were determined separately for the intradepot and peri-depot compartments. Detailed staining and quantification protocols are provided in the Supplementary Information.

### Nonhuman primate studies

Cynomolgus macaques (2.6-4.2 kg, 25-38 months) were randomized into three groups (IV T-DM1, SC TACT alone, SC TACT(T-DM1); n = 3 per group). Animals received 10 mg kg^-1^ T-DM1 either IV or SC as TACT(T-DM1) (650 µL) in the thigh. Plasma T-DM1 and total trastuzumab were quantified by electrochemiluminescence and free DM1 by LC-MS/MS (Eurofins ADME Bioanalyses). Serial ultrasonography with 3D segmentation monitored depot degradation. Full-thickness skin biopsies were collected at day 29. All procedures were approved by CREMEAS (APAFIS #52549-2025010212448791) and performed at SILABE-Université de Strasbourg (Niederhausbergen, France).

### Statistical analysis

Data are presented as mean ± SEM unless otherwise indicated. Pore size distributions, being non-normal, are reported as median [IQR]. Two-group comparisons used two-sided unpaired Mann-Whitney U tests; multiple-group comparisons used Bonferroni-corrected pairwise tests. All statistical analyses were performed in GraphPad Prism. Exact tests and multiple-comparison corrections for each analysis are detailed in the corresponding figure legends.

## Supporting information

Supplementary Informations

## Author contributions

G.J., O.Ti., S.H., C.G., X.P., and A.D. conceived the ideas and designed the experiments. G.J., C.He., E.B., A.G., C.Ho., O.Tr., M.G.-B., J.D., S.E., S.Br., L.F., J.M., E.H., S.Be., J.C., J.G., M.C.A., A.C., and C.M. conducted the experiments. G.J., C.He., E.B., P.C., M.W.T., J.C., L.D., S.C., F.L., O.Ti., S.H., C.G., X.P., and A.D. analyzed the data. S.H., C.G., X.P., and A.D. wrote the manuscript.

## Acknowledgements

We acknowledge Pascal Kessler and Ignacio Busnelli from the PIC’Stra imaging platform. The authors also acknowledge Yacine Ghomari and Anne Rodallec from COMPO team at Institut Paoli Calmettes for their input and assistance with the NCA analyses. We further acknowledge Serena Rizzuti and Eliana Gianolio from Dept. of Molecular Biotechnologies and Health Sciences, Torino, for MRI imaging on mice.

## References

1. Richter, W. F., Bhansali, S. G. & Morris, M. E. Mechanistic determinants of biotherapeutics absorption following SC administration. AAPS J 14, 559–570 (2012).

2. Davis, J. D. et al. Subcutaneous Administration of Monoclonal Antibodies: Pharmacology, Delivery, Immunogenicity, and Learnings From Applications to Clinical Development. Clin Pharmacol Ther 115, 422–439 (2024).

3. Jacquot, G. et al. Landscape of Subcutaneous Administration Strategies for Monoclonal Antibodies in Oncology. Adv Mater 36, e2406604 (2024).

4. Ismael, G. et al. Subcutaneous versus intravenous administration of (neo)adjuvant trastuzumab in patients with HER2-positive, clinical stage I-III breast cancer (HannaH study): a phase 3, open-label, multicentre, randomised trial. Lancet Oncol 13, 869–878 (2012).

5. Pivot, X. et al. Preference for subcutaneous or intravenous administration of trastuzumab in patients with HER2-positive early breast cancer (PrefHer): an open-label randomised study. Lancet Oncol 14, 962–970 (2013).

6. Davies, A. et al. Efficacy and safety of subcutaneous rituximab versus intravenous rituximab for first-line treatment of follicular lymphoma (SABRINA): a randomised, open-label, phase 3 trial. Lancet Haematol 4, e272–e282 (2017).

7. Pivot, X. et al. Patients’ preferences for subcutaneous trastuzumab versus conventional intravenous infusion for the adjuvant treatment of HER2-positive early breast cancer: final analysis of 488 patients in the international, randomized, two-cohort PrefHer study. Ann Oncol 25, 1979–1987 (2014).

8. Locke, K. W., Maneval, D. C. & LaBarre, M. J. ENHANZE® drug delivery technology: a novel approach to subcutaneous administration using recombinant human hyaluronidase PH20. Drug Deliv 26, 98–106 (2019).

9. Bookbinder, L. H. et al. A recombinant human enzyme for enhanced interstitial transport of therapeutics. J Control Release 114, 230–241 (2006).

10. Erfani, A. et al. Crystalline Antibody-Laden Alginate Particles: A Platform for Enabling High Concentration Subcutaneous Delivery of Antibodies. Adv Healthcare Materials 12, 2202370(2023).

11. Erdi, M. et al. High Concentration Antibody Formulations Enabled via Thermostable Ionic Liquids. Advanced Materials 38, e11918 (2026).

12. Schieferstein, J. M., Reichert, P., Narasimhan, C. N., Yang, X. & Doyle, P. S. Hydrogel Microsphere Encapsulation Enhances the Flow Properties of Monoclonal Antibody Crystal Formulations. Advanced Therapeutics 4, 2000216(2021).

13. Drago, J. Z., Modi, S. & Chandarlapaty, S. Unlocking the potential of antibody-drug conjugates for cancer therapy. Nat Rev Clin Oncol 18, 327–344 (2021).

14. Beck, A., Goetsch, L., Dumontet, C. & Corvaïa, N. Strategies and challenges for the next generation of antibody-drug conjugates. Nat Rev Drug Discov 16, 315–337 (2017).

15. Dumontet, C., Reichert, J. M., Senter, P. D., Lambert, J. M. & Beck, A. Antibody-drug conjugates come of age in oncology. Nat Rev Drug Discov 22, 641–661 (2023).

16. Mahalingaiah, P. K. et al. Potential mechanisms of target-independent uptake and toxicity of antibody-drug conjugates. Pharmacology & Therapeutics 200, 110–125 (2019).

17. Detappe, A., Gutiérrez-Blanco, M. & Pivot, X. Engineering the subcutaneous delivery of antibody-drug conjugates. Journal of Controlled Release 395, 114941(2026).

18. Bhattarai, N., Gunn, J. & Zhang, M. Chitosan-based hydrogels for controlled, localized drug delivery. Adv Drug Deliv Rev 62, 83–99 (2010).

19. Gréa, T. et al. Subcutaneous Administration of a Zwitterionic Chitosan-Based Hydrogel for Controlled Spatiotemporal Release of Monoclonal Antibodies. Adv Mater 36, e2308738 (2024).

20. Hennink, W. E. & van Nostrum, C. F. Novel crosslinking methods to design hydrogels. Adv Drug Deliv Rev 54, 13–36 (2002).

21. Kasse, C. M. et al. Subcutaneous delivery of an antibody against SARS-CoV-2 from a supramolecular hydrogel depot. Biomater Sci 11, 2065–2079 (2023).

22. Jons, C. K. et al. Hydrogel formulations for sustained-release of broadly neutralizing antibodies. J Control Release 388, 114349(2025).

23. Verma, S. et al. Trastuzumab emtansine for HER2-positive advanced breast cancer. N Engl J Med 367, 1783–1791 (2012).

24. Lambert, J. M. & Chari, R. V. J. Ado-trastuzumab Emtansine (T-DM1): an antibody-drug conjugate (ADC) for HER2-positive breast cancer. J Med Chem 57, 6949–6964 (2014).

25. Cortés, J. et al. Trastuzumab Deruxtecan versus Trastuzumab Emtansine for Breast Cancer. N Engl J Med 386, 1143–1154 (2022).

26. Modi, S. et al. Trastuzumab Deruxtecan in Previously Treated HER2-Low Advanced Breast Cancer. N Engl J Med 387, 9–20 (2022).

27. Powles, T. et al. Enfortumab Vedotin in Previously Treated Advanced Urothelial Carcinoma. N Engl J Med 384, 1125–1135 (2021).

28. Powles, T. et al. Enfortumab Vedotin and Pembrolizumab in Untreated Advanced Urothelial Cancer. N Engl J Med 390, 875–888 (2024).

29. Ansell, S. M. et al. Overall Survival with Brentuximab Vedotin in Stage III or IV Hodgkin’s Lymphoma. N Engl J Med 387, 310–320 (2022).

30. Korsmeyer, R. W., Gurny, R., Doelker, E., Buri, P. & Peppas, N. A. Mechanisms of solute release from porous hydrophilic polymers. International Journal of Pharmaceutics 15, 25–35 (1983).

31. Krop, I. E. et al. Phase I study of trastuzumab-DM1, an HER2 antibody-drug conjugate, given every 3 weeks to patients with HER2-positive metastatic breast cancer. J Clin Oncol 28, 2698–2704 (2010).

32. Tarantino, P., Ricciuti, B., Pradhan, S. M. & Tolaney, S. M. Optimizing the safety of antibody-drug conjugates for patients with solid tumours. Nat Rev Clin Oncol 20, 558–576 (2023).

33. Tarantino, P. et al. Interstitial Lung Disease Induced by Anti-ERBB2 Antibody-Drug Conjugates: A Review. JAMA Oncol 7, 1873–1881 (2021).

34. Pérez Fidalgo, J. A. et al. Management of chemotherapy extravasation: ESMO-EONS Clinical Practice Guidelines. Ann Oncol 23 Suppl 7, vii167–173 (2012).

35. Kean, T. & Thanou, M. Biodegradation, biodistribution and toxicity of chitosan. Adv Drug Deliv Rev 62, 3–11 (2010).

