## Supplementary Informations for "Hydrogel confinement enables subcutaneous delivery of vesicant antibody-drug conjugates"

2 Equipe labellisée Ligue contre le cancer

3 Strasbourg Drug Discovery and Development Institute (IMS), 67000 Strasbourg, France

4 Nano-H, 69270 Fontaines-Saint-Martin, France

5 Institut Lumière Matière (ILM), Université Lyon 1, CNRS, UMR 5306, Villeurbanne, 69622, Cedex, France

6 Laboratoire de Spectrométrie de Masse BioOrganique (LSMBO), IPHC UMR 7178, Université de Strasbourg, CNRS, 67087 Strasbourg, France; Infrastructure Nationale de Protéomique ProFI - UAR2048, 67087 Strasbourg, France

7 Poly-Dtech, Strasbourg, France

8 SILABE, Université de Strasbourg, Niederhausbergen, France

9 Exercise physiology laboratory, Institut Strauss, Strasbourg; France Biomedicine Research Center of Strasbourg (CRBS), UR 3072, “Mitochondria, oxidative stress and muscle plasticity”; University of Strasbourg, Strasbourg, France European Center of Research, Teaching, Innovation and Exercise Physiology (CEERIPE); Faculty of Sport Sciences, University of Strasbourg, Strasbourg, France

10 Macromolecular Engineering Laboratory, Department of Mechanical and Process Engineering, ETH Zurich, Zurich, Switzerland

11 Université de Strasbourg, Institut Charles Sadron (UPR CNRS 22), Strasbourg Cedex 67034, France

12 Faculté de Médecine de Strasbourg, Institut d'Histologie, Université de Strasbourg, Strasbourg, 67000, France.

13 Ingénierie des Matériaux Polymères (IMP), Université Lyon 1, INSA de Lyon, Université Jean Monnet, CNRS, UMR 5223, Villeurbanne, F69622, Cedex, France

14 Université Bourgogne Europe, Centre Georges-François Leclerc, Unicancer, X-RAIN Unit, Radiation Oncology Department, IMATHERA, UMS INSERM 58 and CTM, UMR INSERM 1231, TIRECs team, 21000 Dijon, France

15 Institut Universitaire de France (IUF), France

16 Recobia Therapeutics, Strasbourg, France

17 Université Paris-Saclay, INSERM, Gustave Roussy, Biothérapies Innovantes U1363, 94800 Villejuif, France

† indicates equal contribution

### Table of Contents

**Section 1. Materials / General Methods / Instrumentation**

**Section 2. Synthetic and formulation procedures**

**Section 3. Supplementary figures**

**Section 4. Supplementary tables**

**Section 5. Supplementary References**

### Section 1. Materials / General Methods / Instrumentation

#### 1.1 Materials

Unless otherwise stated, reagents were obtained from commercial suppliers and used as received. All solutions were prepared using Milli-Q water (18.2 MΩ·cm) and filtered when appropriate.

##### 1.1.1 Polymers and chemicals

Chitosan (Heppe Medical Chitosan; degree of acetylation 2.6 ± 0.6%) served as the starting polysaccharide. The number-average and weight-average molar masses and polydispersity (Mn = 1.362×10^5^ g mol^-1^ ± 2.543%, Mw = 1.995×10^5^ g mol^-1^ ± 0.647%, D = 1.464 ± 2.386%) were determined by HPLC-SEC-MALS (Section 1.2.4). Diacetyl-L-tartaric anhydride (Fischer Scientific, CAS: 6283-74-5, 97%) was used for chitosan functionalization.

Buffers and salts: phosphate-buffered saline (PBS or DPBS; 1X and 5X). Additional salts/acids/bases (HCl, NaOH) were used for pH adjustment.

##### 1.1.2 Antibodies, ADCs and excipients

All antibodies and ADCs were supplied as clinical lyophilized vials by the pharmacy department of the Institut Strauss, Strasbourg, and reconstituted according to manufacturers’ instructions:

• Trastuzumab (Trazimera^®^, Pfizer): used as DAR 0 reference.

• Trastuzumab emtansine (T-DM1, Kadcyla®, Roche; DAR ∼3.5; non-cleavable MCC-DM1 linker).

• Trastuzumab deruxtecan (T-DXd, Enhertu^®^, Daiichi-Sankyo; DAR ∼8; cleavable peptide-DXd linker).

• Enfortumab vedotin (EV, Padcev^®^, Astellas/Pfizer; DAR ∼3.8; cleavable mc-vc-PAB-MMAE linker).

Herceptin SC formulation (Roche) containing recombinant human hyaluronidase (rHuPH20; 2,000 U per vial) was used to prepare dispersion-control groups at a final concentration of 1,000 U after dilution with T-DM1.

Custom trastuzumab-MMAE conjugates (DAR 0, 2, 3.5, and 8; mc-vc-PABC-MMAE architecture) were synthesized in-house (See Section 2.5).

##### 1.1.3 Consumables and sample holders

Needles: 25G hypodermic needles (0.50 × 16 mm, Terumo Agani) for injectability studies and NHP administration; 30G insulin syringes (BD Micro-Fine+ Demi, 0.3 mL) for mouse studies. Syringes: 1 mL Luer-lock syringes for syringe-to-syringe mixing and injectability measurements; BD Hylok glass prefillable syringes (1 mL) for hydrogel sterilization; female-female Luer connector for transfer. Quartz capillaries (Ø 1.5 mm) for SAXS measurements, sealed with epoxy.

#### 1.2 General methods and instrumentation

##### 1.2.1 pH measurement and gelation principle

TACT precursor solutions were prepared at acidic pH (pH 5.7) where the polymer remains a viscous solution. At this pH, the diacetyl-L-tartaric acid moieties retain negatively charged carboxyl groups (estimated pKa ~ 3-4), conferring a polyelectrolytic (zwitterionic) character on the system. At high polymer concentration, the density of intermolecular interactions (hydrogen bonds, hydrophobic interactions and electrostatic contacts between protonated amines and carboxylates) is sufficient to structure a soft hydrogel network within the syringe.

##### 1.2.2 Nuclear magnetic resonance (NMR) spectroscopy

NMR was used to confirm tartrate functionalization and determine the degree of functionalization (DF). ^1^H NMR spectra were acquired on a Bruker Ascend 400 MHz instrument (70°C; 128 scans; pulse program zg30). Samples were prepared at 1% (w/w) in D_2_O; for raw chitosan solubilization, DCl in D_2_O (5% w/w) was used. Spectra were processed in TopSpin (v4.1.3) with phase and baseline correction. DF was determined by integrating the L-tartaric acid methine protons (δ = 5.8 ppm, 2H) relative to the anomeric proton H1 (δ = 5.2 ppm), yielding DF = 25 ± 1% (see Section 2.1.3 for full signal attribution). ^13^C NMR and ^1^H-^13^C HSQC spectra confirmed the assignments. DOSY NMR demonstrated co-alignment of chitosan and L-tartaric acid resonances, supporting a persistent association between the two moieties.

##### 1.2.3 Fourier-transform infrared (FTIR) spectroscopy

FTIR spectra were collected on an IRAffinity-1S spectrophotometer (Shimadzu) equipped with a DLaTGS detector (64 scans, 4 cm^-1^ resolution, 600-4000 cm^-1^ window). Samples were dried at 50°C for 1 h prior to analysis. Atmospheric compensation was applied to minimize CO_2_ and residual water contributions, and a linear baseline correction was applied with the baseline anchored at 4000 cm^-1^.

##### 1.2.4 HPLC-SEC-MALS

Molar-mass and size distributions were determined using a VWR degasser; an Agilent 1260 Infinity pump (0.5 mL min^-1^); an Agilent 1260 Infinity ALS autosampler (100 µL injection); an Agilent 1260 Infinity TCC column oven (30°C); an Agilent 1260 Infinity MWD VL UV detector (280 nm); a Wyatt DAWN HELEOS II MALS detector (662 nm, fused silica cell); and a Wyatt Optilab T-rEX refractive index detector (658 nm, 25°C). Two TSK PW G6000 columns were used in series. Samples were filtered through a 0.45 µm filter prior to injection. The dn/dc value was determined from 0.1-1.0 mg mL^-1^ solutions prepared in mobile phase filtered through a 0.1 µm filter.

##### 1.2.5 SEC-native mass spectrometry (SEC-nMS)

For commercial antibodies: SEC-nMS was used to assess antibody/ADC integrity after gel residence. Samples (10-50 µg) were analyzed on a BioAccord LC-MS system (Waters; ACQUITY UPLC I-Class PLUS coupled to an ACQUITY RDa detector with ESI source and TOF analyzer), controlled via UNIFI. Separation was performed at room temperature on an ACQUITY Premier Protein SEC column (4.6 × 150 mm, 1.7 µm, 250 Å) under isocratic conditions using 150 mM ammonium acetate (pH 6.9). The flow was kept at 150 µL min^-1^ for 0-15.0 min. The RDa detector operated in full-scan positive mode (400-9000 m/z; 2 Hz). Source parameters: capillary 1.50 kV, cone 160 V, desolvation temperature 250 °C. Intact-level samples were injected without additional preparation. For T-DM1, a deglycosylated counterpart was also analyzed to reduce spectral heterogeneity arising from the combination of heterogeneous drug loading and Fc N-glycosylation; deglycosylation was performed by incubating one unit of IgGZERO® enzyme (Genovis, Lund, Sweden) per microgram of ADC for 30 min at 37 °C. For middle-up level analyses, IdeS digestion was performed by incubating one unit of FabRICATOR® enzyme (Genovis, Lund, Sweden) per microgram of ADC for 30 min at 37 °C, generating F(ab')_2_ and Fc fragments for T-DM1, and Fab and Fc fragments for T-DXd.

For T-MMAE ADCs: Native size-exclusion chromatography-mass spectrometry (SEC-MS) was used to characterize antibody-drug conjugate (ADC) species. Analyses were performed using a NativePac OBE-1 column with 50 mM ammonium acetate (pH 6.9) as the mobile phase at a flow rate of 0.1 mL min^-1^. Samples were injected at a concentration of 1-2 mg mL^-1^, with a total acquisition time of 5 minutes. The column eluent was directly introduced into an Exploris 480 mass spectrometer equipped with an electrospray ionization (ESI) source operating in positive ion mode at a resolution of 30,000 under native conditions. Full MS spectra were acquired over an m/z range of 3000*6000. Data were acquired using Xcalibur 4.7 software, processed with Freestyle 1.8, and deconvoluted with UniDec to obtain the molecular masses of the detected species and the drug-to-antibody ratio (DAR) distribution.

Average DAR values were determined from the relative intensities of deconvoluted DAR species using the weighted-average DAR (equation 1). Representative spectra are shown in Supplementary Fig. 7 and the corresponding DAR distributions are summarized in Supplementary Table 10.

$\text{DAR=}\frac{\left( \sum_{\text{n=0}}^{\text{8}} \text{n*intensity DARn} \right)}{\sum_{\text{k=0}}^{\text{8}} \text{intensity DARn}}$ **(eq. 1)**

##### 1.2.6 SEC analysis of T-MMAE ADCs aggregation

The aggregation state of T-MMAE antibody-drug conjugates (ADCs) was evaluated by size-exclusion chromatography (SEC) using an ÄKTA FPLC system (Cytiva) fitted with a Superdex 200 Increase 10/300 GL column. Samples (50 µL at 1 mg mL^-1^) were injected through a sample loop and eluted with PBS as the mobile phase at a flow rate of 0.5 mL min^-1^. Protein elution was monitored by UV absorbance at 280 nm. The relative percentage of high-molecular-weight (HMW) species, monomer, and low-molecular-weight (LMW) fragments was determined by integration of the chromatographic peaks and expressed as a percentage of the total peak area. (Supplementary Fig. 6)

##### 1.2.7 Rheology

Dynamic shear rheology was performed on a strain-controlled rheometer (Discovery HR-10, TA Instruments) equipped with a 20 mm sand-blasted stainless-steel parallel-plate geometry and Peltier temperature control at 25 °C. A gap of 0.5 mm was used for all measurements. Approximately 150 µL of hydrogel was loaded from syringes and an oscillatory pre-conditioning step (ω = 1 rad s^-1^, γ = 0.1%, 3 min) was applied to standardize loading history. Strain sweeps (0.1-1000% at ω = 10 rad s^-1^) defined the linear viscoelastic range. Frequency sweeps were recorded from 100 to 0.1 rad s^-1^ at 1% strain. Flow behavior was characterized by shear-rate ramps from 0.1 to 100 s^-1^. Self-healing properties were characterized with oscillatory time sweep measurements with alternating 5-min intervals of low strain (γ = 1% and ω = 10 rad s^−1^) and high strain (γ = 1000% and ω = 10 rad s^-1^). All measurements were performed in duplicate.

##### 1.2.8 Injectability and extrusion force measurements

Injectability was assessed on a PHD 22/2000 syringe pump mounted with a piezo electrical device with a sensitivity ranging from mN to 200N (FC2231-0000-0050-L). 1 mL luer lock syringe (Aesthetic Group; low dead space) fitted with Terumo Agani 25G needles (0.50 × 16 mm) were secured in a custom chamber with axial polyamide screws. Injection flow rate was set to 1 ml min^-1^. The reported injection force corresponds to the mean force over a 5 s window after reaching a stable plateau. Two test configurations were used: (i) extrusion into air, to isolate syringe-and-needle resistance, and (ii) extrusion into full-thickness human skin explants, to capture tissue backpressure. Tissue backpressure was calculated as ΔP = P_skin_ - P_air_. H&E staining assessed acute tissue architecture after injection.

##### 1.2.9 Cryogenic scanning electron microscopy (cryo-SEM)

The structure of the TACT native hydrogel (2.5% w/w) and ADC-loaded hydrogels (2.5% w/w polymer; T-DXd 50 mg mL^-1^, T-DM1 50 mg mL^-1^) was analyzed under two conditions: intact TACT hydrogels and hydrogels after injection of 200 µL into 1X PBS for 1 h. Subsequently, a small piece of gel was quickly immersed into liquid ethane, then placed under high vacuum (10^-6^ mbar) and low temperature (-150 °C) in the cryo-preparation chamber (Quorum PT 3010) attached to the microscope. There, an adapted razor blade was used to fracture the frozen sample. To reveal morphological details, the material was subjected to two sublimation cycles (3 min, -90 °C; 5 min, -90 °C). Eventually, the sample was transferred to the Field Emission Gun-cryo-SEM (Hitachi)

Pore morphology and distribution were quantified using a custom Python pipeline. Images were acquired at multiple magnifications (5k×-40k×, pixel sizes 19.8-2.5 nm). Pores were segmented using Gaussian-weighted adaptive local thresholding to accommodate heterogeneous illumination across the field of view. This algorithm computes pixel-specific thresholds based on the Gaussian-weighted mean intensity within a 101×101 pixel neighborhood, subtracting a constant offset (C=2):

*Binary(x,y) = 1 if I(x,y) < T_local(x,y) - C, else 0*

Connected components were identified using 8-connectivity and filtered by size (10-5000 pixels^2^, corresponding to ∼5-1000 nm diameter) and shape (circularity > 0.6, solidity > 0.7). For each pore, area, equivalent diameter (D = 2√(A/π)), perimeter, Feret diameters, circularity (4πA/P^2^), eccentricity, aspect ratio and solidity were measured. Images were grouped by magnification for valid comparisons; comparative analyses used only images at identical magnification (typically 20k× or 40k×). Parametric data were analyzed using one-way analysis of variance (ANOVA) followed by Tukey's honestly significant difference post hoc test for pairwise comparisons. Non-parametric distributions (e.g., diameter) were evaluated using the Kruskal-Wallis test. Categorical data (size class distributions) were compared using chi-square tests. Statistical significance was defined as P < 0.05. Data are presented as mean ± standard deviation for continuous variables and median with interquartile range for non-normally distributed parameters. Analytical reproducibility was assessed through coefficient of variation (CV) analysis. Within-formulation CV for pore density ranged from 3.1% to 7.4%, demonstrating excellent sample homogeneity and measurement precision. The fully automated nature of the analysis pipeline eliminated inter-operator variability, ensuring complete reproducibility. Segmentation accuracy was validated against manual measurements on a subset of 50 pores, yielding 94.2% sensitivity and 95.1% positive predictive value.

##### 1.2.10 Small-angle X-ray scattering (SAXS)

Two SAXS datasets were acquired. Dataset 1 (0-17 h, four conditions: TACT alone, TACT(trastuzumab), TACT(T-DM1), TACT(T-DXd)) was collected on a Xeuss SAXS instrument illuminated with Cu Kα radiation (λ = 1.542 Å) at a sample-to-detector distance of 600 mm, covering q = 0.005-0.40 Å^-1^ (real-space length scales D = 2π/q from 1.4 to 126 nm). Exposure time was 900 s per frame; samples were mounted in 1.008 mm-thick capillaries. Dataset 2 (0-98 h, three conditions: TACT alone, TACT(trastuzumab), TACT(T-DM1)) was collected on the DiffériX platform (Institut Charles Sadron, Strasbourg) using a Rigaku Micromax-007 HF rotating anode generator (40 kV, 30 mA; Cu Kα, λ = 1.54 Å) equipped with Max-Flux confocal multilayer optics and a 3-pinhole collimation system, with a 2D multiwire detector at 81 cm (q ∼ 0.01-1.0 Å^-1^). Samples were loaded in sealed quartz capillaries (Ø 1.5 mm) at 25 °C. Silver behenate was used for q-calibration and Lupolen for absolute intensity calibration. All SAXS data reported in the main text refer to Dataset 1 (Xeuss). Extended longitudinal SAXS data (Dataset 2, up to 98 h) are provided in Supplementary Fig. S3.

In the gel state, SAXS probes both the large-scale organization of the polymer network at low q and the local conformational properties of the chains at higher q.

The scattering profiles display a pronounced upturn at very low q (below 0.01 Å^-1^ not shown), characteristic of large-scale heterogeneities within the gel network, together with an overall q^−2^ decay in the asymptotic regime, consistent with Gaussian statistics of the chitosan chains. Similar behaviors are commonly observed in physically cross-linked polymer hydrogels^1^ .

Structural differences are nevertheless observed depending on the nature of the antibody incorporated into the gel. To better reveal these variations, the data were plotted in Kratky representation (q^2^I(q) versus q), which removes the dominant q^−2^ decay and enhances subtle behaviors after subtraction of buffer scattering. In such a representation, Gaussian chains are expected to show a characteristic plateau at high q.

For the pure TACT hydrogel, we observe a simple broad plateau that gradually increases with time. This behavior suggests a slow reorganization and densification of the network while preserving an overall Gaussian-chain network structure, even after 17 h of aging. No clear Lorentzian-like function associated to the mesh-scale organization is visible in Log-Log plots^1^. Only a very small signature is revealed in Kratky representation for the first hours of aging.

In contrast, the presence of antibodies significantly modifies the shape of the scattering curves, indicating a clear reorganization of the polymer network. Although the global q^−2^ dependence remains preserved, two broad maxima located at q_1_ and q_2_ progressively emerge and evolve with time (Kratky plots). This effect is particularly pronounced for trastuzumab-containing gels, for which the SAXS profiles display two marked “shoulders” at the end of the kinetic sequence (Log-Log representation)

These correlation peaks depart from the simple Ornstein-Zernike contribution expected for semidilute Gaussian chains and instead reveal the existence of two characteristic structural length scales within the gel^2^. The first one, observed at lower q, is attributed to correlations between antibody-induced aggregates. In this picture, antibodies act as transient cores onto which chitosan-tartrate chains associate, leading to the formation of star-like aggregates that act as physical cross-linking nodes. The corresponding average inter-aggregate distance, $\Theta$, is related to the position of the first maximum through:

$$\Theta=\frac{2\pi}{q_{1}}$$

The second, broader maximum observed at larger q is assigned to the polymer correlation length $\xi$, corresponding to the average interchain distance within the semidilute polymer network:

$$\xi=\frac{2\pi}{q_{2}}$$

The kinetic evolution observed in antibody-containing gels mainly originates from changes in the position (and shape) of the first correlation peak (q_1_). This suggests that the structural rearrangement is governed by the equilibration dynamics of the antibodies within the network, and therefore, by their interaction capacity with the polymer chains. Over very long aging times (several weeks), the shoulders observed for trastuzumab-containing gels almost completely disappear, indicating a progressive redistribution of the antibodies within the gel.

##### 1.2.11 In vitro release assays for commercial mAbs and ADCs

Simulated SC medium was prepared by dissolving in 1 L of deionized water: NaCl (8.45 g), KCl (0.37 g), CaCl_2_·2H_2_O (0.22 g), MgCl_2_·6H_2_O (0.142 g), NaHCO_3_ (2.10 g), Na_2_HPO_4_·H_2_O (0.142 g), NaH_2_PO_4_·H_2_O (0.069 g) and D-glucose (0.9 g). The pH was adjusted to 7.4 using 1 M HCl and the solution was equilibrated at 25 °C. Release assays were performed at 25 °C (room temperature) for experimental consistency across all time points.

TACT or control hydrogels loaded with antibody or ADC (100 µL) were injected into 1.5 mL tubes and 1 mL of simulated SC medium was added. At predefined time points (0.25-6 h), samples were gently homogenized and 2 µL of supernatant were withdrawn in triplicate. Concentrations were quantified by UV absorbance at 280 nm (NanoPhotometer, Implen). Percentage of release was calculated as the ratio between measured antibody concentration and theoretical total antibody concentration (based on hydrogel volume added). Release from unmodified chitosan was measured under identical conditions (n = 3). Ionic-strength dependence was assessed by substituting SC medium with water, 1X DPBS or 5X DPBS (n = 2-3 per condition).

##### 1.2.12 In vitro release assays for Trastuzumab-VC-MMAE

Release assays were conducted by adding ≈100 μL of TACT (2.5% w/w) hydrogel loaded with antibody or ADC (final concentration: 11.25 mg mL^-1^) to an Eppendorf tube (1.5 mL). Samples were briefly centrifuged (a few seconds) until the hydrogel was situated at the bottom of the tube. The experiment was started by carefully adding 1 mL of 1X PBS on top of the hydrogel and incubating the samples at room temperature. At predefined time points (0.25-6 h), the supernatant was gently homogenized and 50 μL of the sample was collected and replaced with fresh PBS. Antibody concentrations were quantified by UV absorbance at 280 nm. All the experiments were repeated in triplicate. The percentage of release was calculated as the ratio between measured antibody concentration (equation 2) and theoretical total antibody concentration (based on hydrogel volume added). To account for the removal of antibody at each sampling point, the cumulative mass of antibody removed in previous aliquots was included in the calculation

m_t_ = C_t_·V +$\text{∑}_{\text{i=1}}^{\text{t-1}}$C_i_·V_s_	(**eq.2**)

where:

m_t_ = cumulative mass of antibody release at time t

C_t_ = antibody concentration measured at time t

V = total volume of the release medium

C_i_ = antibody concentration measured at the previous sampling points

V_s_ = sample volume withdrawn at each time point.

##### 1.2.13. In-vitro binding assays

Flow cytometry data acquisition was performed using the MACSQuant Analyzer 10 Flow Cytometer (Miltenyi), and subsequent analysis was carried out using FlowJo software (V. 10.8.2).

HCC1954 cells were seeded in a 96-well plate at a density of 5 × 10^4^ cells per well and pelleted by centrifugation at 300 × g for 5 min. Supernatants were carefully removed, and cells were washed once with Dulbecco’s phosphate-buffered saline (DPBS).

Serial dilutions (1:2) of unlabeled antibody or antibody-drug conjugate (ADC) were prepared in DPBS, starting from 1 mg mL^-1^. Cells were incubated with 100 µL of the corresponding antibody dilution for 10 min at 4 °C. Following incubation, cells were centrifuged (300 × g, 5 min) and washed once with 100 µL of DPBS to remove unbound antibody. Cells were then incubated with 100 µL of fluorescently labeled trastuzumab at a final concentration of 0.01 mg mL^-1^ (previously optimized by titration) for 10 min at 4 °C. After incubation, cells were centrifuged (300 × g, 5 min) and washed three times with 200 µL of DPBS, with centrifugation between each wash step. Finally, cells were resuspended in 200 µL of DPBS and analyzed by flow cytometry.

##### 1.2.14 Pharmacokinetics in mice

All mouse experiments were approved by the relevant institutional animal care and use committee.

T-DM1: Female adult C57BL/6J mice (16-25 g) were used. TACT hydrogels at 2.5% w/w were mixed with T-DM1 to obtain 50 mg mL^-1^. Mice were separated into three groups (n = 19 per group): IV T-DM1 17.5 mg kg^-1^, SC TACT(T-DM1) 17.5 mg kg^-1^ (7 µL) or 30 mg kg^-1^ (12 µL). Study duration: 42 days.

T-DXd: Female adult C57BL/6J mice (16-25 g). TACT at 2.5% w/w, 50 mg mL^-1^. Two groups (n = 19 per group): IV T-DXd 10 mg kg^-1^ or SC TACT(T-DXd) 10 mg kg^-1^ (4 µL). Study duration: 42 days.

Trastuzumab: Female adult BALB/c mice (16-25 g). TACT at 5% w/w, 100 mg mL^-1^. Three groups: IV trastuzumab 500 mg kg^-1^ (100 µL, n = 9), SC Herceptin 600 mg kg^-1^ (100 µL, n = 9), or TACT(trastuzumab) 500 mg kg^-1^ (100 µL, n = 16). Study duration: 28 days.

For IV groups, tail-vein injection was performed. SC injections were performed in the dorsal flank using 30G insulin needles (for volumes up to 12 µL) or using 1 mL luer lock syringe combined with 25G hypodermic needles (for 100 µL injection volume). At each sampling time point, blood was collected from a subset of animals via the mandibular vein into EDTA-coated tubes, centrifuged, and plasma stored at -20 °C. For selected time points, mice were euthanized and the injection site excised for histological analysis. Plasma concentrations were quantified using a TR-FRET assay (HTRF Human IgG Detection kit, Cisbio/Revvity) or TR-FLISA assay (Poly-Dtech kit). Noncompartmental pharmacokinetic parameters were derived from mean concentration-time profiles.

**1.2.15. Trastuzumab quantification by Time-resolved fluorescence energy transfer (TR-FRET)**.

HTRF Human IgG Detection kit (Cisbio, Revvity) is a sandwich immunoassay based on Homogenous Time-Resolved Fluorescence (HTRF). Plasma trastuzumab level were measured in mice according to the manufacturers’ instructions. Plasma samples were diluted in assay buffer at optimized dilution factors (from 500 to 10000-fold) and analyzed in triplicate. The calibration curve was obtained by serial dilution of a trastuzumab stock solution (Trazimera, Pfizer) in assay buffer to cover the range of 0-2000 ng mL^-1^, assay buffer alone was used as a negative control. Ten µL of standards and samples were then dispensed into a 384-well plate, followed by the addition of 10 µL of the detection reagent mixture provided in the kit. Plates were incubated for 3h, at room temperature (RT), protected from the light. Finally, fluorescence was subsequently read at 615 nm and 665 nm (excitation at 340 nm) using the TR-FRET mode of a SpectraMax ID5 plate reader (Molecular Devices), with settings configured according to the Cisbio manufacturer’s guidelines. Raw data were processed using SoftMax Pro software.

**1.2.16. Time resolved-fluorescence linked immunosorbent assay (TR-FLISA).**

TR-FLISA is a variant of the enzyme-linked immunosorbent assay (ELISA) technique which replaces the enzymatic reaction by time-resolved fluorescent molecules. Plasma levels of the ADCs (T-DM1, T-DXd) were measured using the mice TR-FLISA kits from Poly-Dtech (Strasbourg, France, FLI-MouTRASTU) and assays were performed according to the manufacturers’ instructions. In brief, plates were washed 3 times with washing buffer. Then, 100 µL of standards and samples were added in duplicate. An ADC of interest standard curve was used for each assay. Dilution buffer was used as the zero standard (0 ng mL^-1^) and plasma from mice before administration was used as negative control. Different plasma dilutions were tested for all samples to establish the experimental setup. Plates were incubated with gentle shaking for 1h at 37 °C followed by 3-times washing. After what, 100 µL of the detection antibody, diluted in assay buffer, was added to each well and incubated with shaking for 1h at 37°C. For the final wash, the plates were washed 6 times (5 minutes incubation with shaking per wash) to help minimize background. Then, 200 µL of washing buffer was added to each well and fluorescence was measured at 535 nm (excitation at 340 nm) using the time resolved fluorescence (TRF) read mode of a SpectraMax ID5 (Molecular Devices) following the TRF plate reader settings in the manufacturer’s instructions. Raw data were processed with the corresponding SoftMax Pro software using a 5-parameter logistic curve.

##### 1.2.17. Xenograft efficacy studies

BT-474 cells were grown as monolayers at 37 °C in humidified atmosphere (5% CO_2_, 95% air) in DMEM containing 2 mM L-glutamine and 1 mM sodium pyruvate supplemented with 10% fetal bovine serum. Cells were detached by 5 min treatment with trypsin-versene and neutralized with complete medium.

BT-474 cells (2×10^7^) in 50% (v/v) Matrigel were implanted SC in female athymic nude mice (n = 8 per group) and animals received water supplemented with 2.5 µg mL^-1^ estradiol for the duration of the study. Treatment began at ∼160 mm^3^.

For T-DM1: TACT hydrogels at 2.5% w/w were mixed with T-DM1 to obtain 50 mg mL^-1^. For T-DM1 single-dose: IV Kadcyla (17.5 mg kg^-1^) versus SC TACT(T-DM1) (17.5 or 30 mg kg^-1^). For T-DM1 multi-dose: Q7D×4 at 5 mg kg^-1^.

For T-DXd single-dose: TACT hydrogels at 2.5% w/w were mixed with T-DXd to obtain 50 mg mL^-1^, IV T-DXd (10 mg kg^-1^) versus SC TACT(T-DXd) (10 mg kg^-1^).

For trastuzumab: TACT hydrogels at 5% w/w were mixed with Trastuzumab to obtain 100 mg mL^-1^, Q7D×4 of IV Trazimera (10 mg kg^-1^), SC Herceptin (15 mg kg^-1^) or SC TACT(trastuzumab) (15 or 30 mg kg^-1^). All studies included PBS and TACT alone control groups. Body weights and tumor volumes (length × width^2^/2) were measured at least twice weekly.

##### 1.2.18. Local tolerance, histology and immunohistochemistry

For local tolerance, C57BL/6 mice received SC injections of 30 mg kg^-1^ T-DM1 (Kadcyla), TACT(T-DM1), T-DM1 + rHuPH20 (1,000 U) or TACT alone. Because formulation concentrations differed, injection volumes were not identical; for a 20 g mouse, approximate administered volumes were 12 µL for TACT(T-DM1), 60 µL for T-DM1 alone and 60 µL for T-DM1 + rHuPH20. These volume differences were recorded and considered during interpretation of local inflammation and necrosis; the rHuPH20 group was used as a dispersion-control formulation rather than a volume-matched control. Sites were inspected daily. All biopsies were collected from the flank at the injection site at predetermined sacrifice time points. Excised tissues were fixed in 4% paraformaldehyde for 24 h, dehydrated through graded ethanol solutions, cleared in toluene, and embedded in liquid paraffin for 24 h.

Serial 5 µm sections were used for H&E, anti-human Fc/anti-trastuzumab immunohistochemistry, TUNEL labeling, and F4/80 and Ly6G immunostaining. TUNEL labeling was performed on deparaffinized sections according to the manufacturer's instructions. F4/80 and Ly6G staining were performed after antigen retrieval, blocking, primary antibody incubation, chromogenic detection and hematoxylin counterstaining. Whole-slide images were analyzed by automated thresholding to quantify TUNEL+ tissue area and by compartmental cell-density measurements for F4/80+ and Ly6G+ cells in intradepot and peri-depot regions.

Sections (5 µm) were deparaffinized and rehydrated through sequential toluene, ethanol and distilled water baths. Hematoxylin and eosin (H&E) staining was performed (2 min each). After staining, sections were dehydrated and mounted using Eukitt®.

##### 1.2.19. Non-human primate (NHP) pharmacokinetics and safety

Cynomolgus macaques (2.6-4.2 kg, 25-38 months) were randomized into three groups (n = 3 per group): IV T-DM1, SC TACT alone and SC TACT(T-DM1). Animals were anesthetized with ketamine (10 mg kg^-1^) and medetomidine (Domitor, 2 mg kg^-1^) by intramuscular injection. Treated groups received 10 mg kg^-1^ T-DM1 either intravenously or subcutaneously as TACT(T-DM1) (650 µL of gel) in the thigh. The TACT alone group received the same gel volume.

At each time point, animals were anesthetized as above for ultrasound imaging and blood collection. Plasma sampling times: T0, 6 h, 12 h, D1, D2, D3, D4, D5, D6, D9, D11, D16, D18, D23, D25 post-treatment. Depot monitoring was performed by 2D ultrasonography (General Electric Versana Active 1.5, 12L-RS linear probe, 12.0 MHz). Five images were acquired per time point and analyzed by three independent investigators in a blinded manner using ImageJ (v1.8.0).

Plasma T-DM1 and total trastuzumab were quantified by electrochemiluminescence: quantification range of 2-100 ng mL^-1^ and 0.4-25 ng mL^-1^ respectively (samples diluted 100-20,000-fold). Free DM1 was quantified by LC-MS/MS (quantification range: 0.1-30 ng mL^-1^; samples analyzed without dilution). All analyses were performed by Eurofins ADME Bioanalyses. DM1 catabolite fraction was calculated as the ratio of measured free DM1 to total theoretical DM1 concentration, derived from: C_theoretical_, DM1 = (C_T-DM1_ / MW_T-DM1_) × MW_DM1_ × DAR, where MW_T-DM1_ = 147,781 g mol^-1^, MW_DM1_ = 738.8 g mol^-1^ and DAR = 3.5.

Full-thickness skin biopsies (8 mm punch) were collected at day 26 and fixed in 4% PFA for 24 h. Cutaneous reactions at SC injection sites were observed daily. Animal procedures were approved by CREMEAS (APAFIS #52549-2025010212448791) and performed at SILABE-Université de Strasbourg (Niederhausbergen, France).

#### 1.3 Data analysis and software

Data are presented as mean ± SEM unless otherwise indicated. Non-normal distributions are reported as median [IQR]. Two-group comparisons used two-sided unpaired Mann-Whitney U tests. Multiple-group comparisons used Kruskal-Wallis tests with Dunn’s post hoc correction, except for cryo-SEM pore-size comparisons which used Bonferroni correction (six pairwise tests; adjusted threshold P < 0.0083). The exact statistical test and correction applied for each primary comparison are specified in the corresponding figure legend. The IVIVC correlation (r = 0.997, p = 0.046) is based on n = 3 cargo conditions; this sample size is insufficient to establish a quantitative IVIVC relationship and should be interpreted as a preliminary correlation pending validation with additional cargo classes. PK analyses were performed in IGOR Pro (v9.0, Wavemetrics) and custom Python scripts. Release kinetic modeling (Korsmeyer-Peppas, Weibull, first-order, Higuchi, Hixson-Crowell, zero-order) used scipy.optimize.curve_fit with model selection by AIC. Immunoassay curve fits were performed in SoftMax Pro (Molecular Devices) using a five-parameter logistic model. Image processing used ImageJ (v1.8.0) and/or Fiji (v2.9.0).

#### 1.4 Animal models, ethics and study design

All animal experiments were approved by the relevant institutional animal care and use committee and conducted in accordance with applicable guidelines.

Animals were randomized to treatment groups by body weight stratification or tumor size. Sample sizes were based on prior experience with similar ADC SC delivery studies and are described per experiment in the relevant Methods subsection. Exclusion criteria: animals showing signs of distress unrelated to treatment were excluded; no exclusions were made based on outcome data.

### Section 2. Synthetic and Formulation Procedures

#### 2.1 Synthesis of TACT polymer

TACT was synthesized in a one-pot procedure by functionalization of diacetyl-L-tartaric anhydride onto chitosan in a water/propanediol mixture at room temperature (Fig. 1a and Supplementary Fig. 1a).

##### 2.1.1 Reaction setup

Chitosan (7.5 g) was dispersed in 500 mL of Milli-Q water under vigorous agitation. Acetic acid (10 mL, 99.5%) was added and the mixture was stirred for 4 h until complete dissolution (expected pH ∼3.1). Propanediol (500 mL, 1,2-propanediol) was then introduced and stirred for 30 min. Diacetyl-L-tartaric anhydride (3.30 g, 0.33 molar equivalents relative to chitosan amines) was added and the reaction proceeded overnight at room temperature. The reaction mixture was filtered through a Büchner funnel (12-15 µm pore size).

##### 2.1.2 Work-up and purification

The product was diluted 4-fold with water and purified by tangential-flow filtration (Sartorius Sartoflow Smart TFF system; 2 bar inlet, 1.5 bar transmembrane pressure; two 0.02 m^2^ cassettes, 50 kDa MWCO). Purification consisted of five cycles with 0.1 M acetic acid followed by five cycles with 5 mM acetic acid, each time reconcentrating the retentate to 250 mL (total: 30 diavolumes; theoretical purification factor ∼10^6^). pH before TFF was 3.8 and reached 5.0 after purification. The purified polymer was freeze-dried to yield TACT as a lyophilisate (pH 5.7 after redispersion at 10% w/w).

##### 2.1.3 Polymer characterization

Functionalization was confirmed by ^1^H NMR spectroscopy in D_2_O at 70°C (Supplementary Fig. 1b). The TACT spectrum exhibited signals from both the chitosan backbone and the diacetyl-L-tartaric moieties, demonstrating that the molecule remains associated with the polymer even after 10^6^-fold purification by TFF. Key signal assignments: δ = 5.80 ppm (Hb, methine between carboxylic acid and ester groups), δ = 5.19 ppm (H1, anomeric proton), δ = 4.0-4.5 ppm (H2-H7, glucosamine ring), δ = 3.47 ppm (H2(D), α to amine in deacetylated units), δ = 2.56 ppm (Ha, methyl protons of L-tartaric unit). Residual water and acetic acid peak were observed at 4.7 ppm and 2.4 ppm respectively. See Supplementary Fig. 1b for annotated structure and full attribution.

^13^C NMR and HSQC confirmed assignments (Supplementary Fig. 1c,d). DOSY NMR showed co-alignment of chitosan and tartrate diffusion coefficients, supporting formation of a single macromolecular species (Supplementary Fig. 1g).

FTIR showed characteristic amide stretching (ν-C=ONH, ∼1604 cm^-1^), ester/carboxylic acid stretching (ν-C=O, ∼1728 cm^-1^) and ester C-O stretching (ν-C-O, ∼1249 cm^-1^) consistent with functionalization (Supplementary Fig. 1e).

The degree of functionalization was determined by ^1^H NMR using: DF = I(5.80 ppm)/2 ÷ I(5.19 ppm) = 25 ± 1%.

HPLC-SEC-MALS yielded Mn = 1.346×10^5^ g mol^-1^ (±1.410%), Mw = 1.959×10^5^ g mol^-1^ (±0.585%), Mp = 1.543×10^5^ g mol^-1^ (±0.261%) and Đ = 1.456 (±1.527%) (Supplementary Fig. 1f).

#### 2.2 Preparation of TACT precursor solutions and hydrogel formation

##### 2.2.1 Precursor solution preparation

Lyophilized TACT was dissolved at 10% (w/v) in Milli-Q water (final pH 5.7). The viscous solution was transferred to Luer-lock syringes, centrifuged (3,500 rpm, 5 min) to remove air bubbles, transferred to autoclavable glass syringes and sterilized by autoclaving (121 °C, 20 min). Sterilization resulted in increased viscosity (from ∼120 to ∼1200 Pa s at 1 s^-1^), consistent with enhanced intermolecular interactions; values remained compatible with injectable administration. Precursor solutions were stored at 4 °C. Stability studies at 4 °C and under accelerated conditions (room temperature and 30 °C) over 1 month showed no degradation.

##### 2.2.2 Gelation in physiological conditions

#### At its pre-injection pH of 5.7, the precursor solution forms an initial polyelectrolyte complex via electrostatic interactions between the protonated amines of the chitosan backbone and the carboxylate groups of the diacetyl-L-tartaric moieties. When TACT is exposed to physiological buffer, gelation proceeds through two stages. First, ion diffusion from the environment screens cationic charges on the chitosan chains, reducing electrostatic repulsion and enabling closer contact between protonated amines and anionic carboxylate groups. Second, as pH reaches 7.4, the majority of chitosan amines are deprotonated, increasing chain hydrophobicity and enabling organization through hydrophobic interactions between glucosamine/N-acetylglucosamine motifs and interchain hydrogen bonds. This two-step mechanism, which distinguishes TACT from unmodified chitosan that precipitates rather than gels at physiological pH, produces a stable, self-healing hydrogel network.

#### 2.3 Loading of antibodies and ADCs

Clinical lyophilized vials were reconstituted at 100 or 200 mg mL^-1^ in water for injection. To prepare the lead formulation (2.5% w/w TACT, 50 mg mL^-1^ ADC), TACT precursor (10% w/w) was first diluted twofold by syringe-to-syringe transfer with water for injection (∼200 reciprocations, 2 min). Equal volumes of diluted precursor (5% w/w) and antibody/ADC solution were then mixed by syringe-to-syringe transfer (∼200 reciprocations, 2 min). Visual inspection confirmed absence of precipitation or phase separation. Formulations at 5% w/w polymer (100 mg mL^-1^ antibody) omitted the preliminary dilution step. Formulations were transferred to BD Hylok syringes (25G) or insulin syringes (30G) and used immediately or stored at 4 °C for up to 1 week. For MMAE-based ADCs, TACT precursor (10% w/w, 100 µL) and antibody solution (15 mg mL^-1^, 300 µL) were mixed by syringe-to-syringe transfer (∼200 reciprocations, 2 min) to obtain a final formulation comprising 2.5% w/w TACT and 11.25 mg mL^-1^ antibody, used immediately for release tests.

#### 2.4 Control formulations

##### 2.4.1 Unmodified chitosan hydrogel

Unmodified chitosan was dissolved in acetic acid (1% v/v) and purified by TFF following the same program as TACT. After freeze-drying, the polymer was dissolved at 10% (w/w) in Milli-Q water (pH 5.0), transferred to syringes, centrifuged and autoclaved. Formulations were prepared at 2.5% w/w with or without 50 mg mL^-1^ trastuzumab by the same syringe-to-syringe protocol. Rheological characterization confirmed that unmodified chitosan formulation is a solution (G’ < G’’) compared to TACT (Supplementary Fig. 5a). Upon immersion in SC medium at pH 7.4, no gel formation was observed, leading to burst release of trastuzumab (Supplementary Fig. 5b).

##### 2.4.2 rHuPH20 co-formulation control

For dispersion-enhancing controls, T-DM1 was co-formulated with rHuPH20 immediately prior to SC injection. One volume of T-DM1 clinical solution (20 mg mL^-1^) was mixed with one volume of Herceptin SC formulation (120 mg mL^-1^ trastuzumab, 2,000 U rHuPH20), yielding a final T-DM1 content of 10 mg mL^-1^ and 1,000 U rHuPH20. Formulation clarity was verified before injection to rule out gross aggregation.

**2.5.** **Synthesis and characterization of Trastuzumab-VC-MMAE**

Trastuzumab-VC-MMAE ADCs were synthesized by conjugating trastuzumab (T) (Trazimera, Pfizer) with the MC-VC-PAB-MMAE drug linker. Three different drug-to-antibody (DAR) ratios were targeted: 2, 4 and 8. For DAR=2, the site-selective GeminiMab technology by Syndivia was employed. For DAR=4 and DAR=8, a conventional random cysteine conjugation method was used.^3^

Trastuzumab (8.45 mg mL^-1^) was partially (2.5 eq.) or fully reduced (8 eq.) with TCEP (tris(2-carboxymethyl) phosphine) (10 mM in water) and incubated at 37 °C for 1.5 h. Next, the solution was cooled down to 4 °C and an excess of MC-VC-PAB-MMAE (10 mM in DMSO) was added. The reaction mixture was incubated for an additional 1.5 h to allow conjugation. The resulting antibody solution was centrifuged at 13,200 g for 4 minutes to remove the insoluble excess of payload followed by purification using Bio-Spin P-30 Gel columns (4 min, 1000 rpm, pre-equilibrated with PBS 1x) to yield purified T-MMAE. The concentration of the ADCs was determined by NanoDrop UV-Vis spectrophotometry. Samples were diluted to 1-2 mg mL^-1^ for chromatographic analysis. The DAR distribution was determined by size-exclusion chromatography coupled to native mass spectrometry (SEC-nMS), while the aggregation state of the conjugates was assessed by SEC using an ÄKTA FPLC system. (Supplementary Figure 6 and 7, Supplementary Table 10).

### Section 3. Supplementary Figures

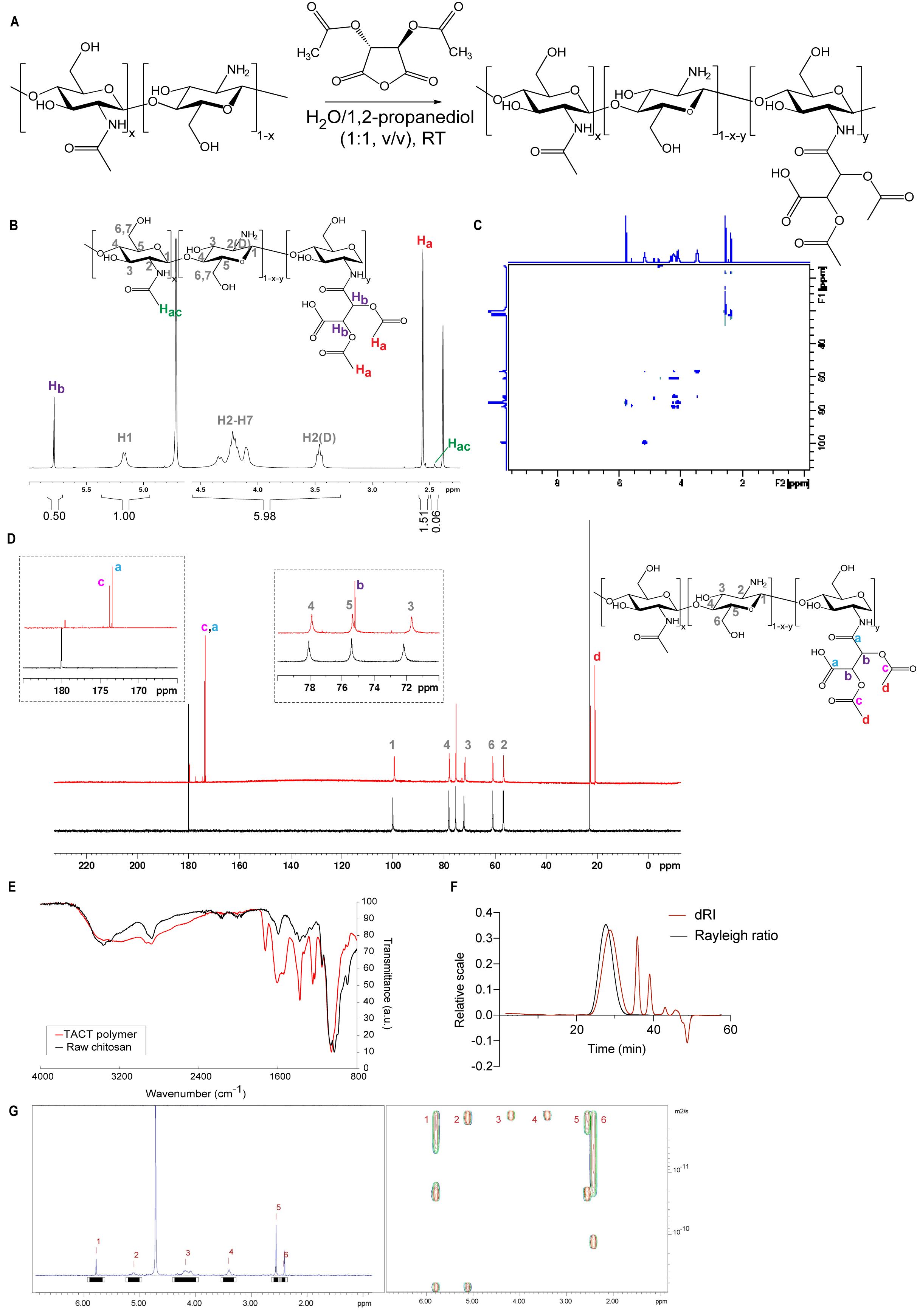

**Supplementary Figure 1.** Synthesis and characterization of TACT polymer. **A.** Reaction schematic. **B.** Representative ^1^H NMR spectra TACT and annotated TACT structure for signal attribution. **C.** ^1^H-^13^C HSQC NMR. **D.** ^13^C NMR spectra of chitosan (black) and TACT polymer (red) and annotated TACT structure for signal attribution. **E.** FTIR spectra of chitosan (black) and TACT (red). **F.** HPLC-SEC-MALS chromatogram. **G.** DOSY NMR.

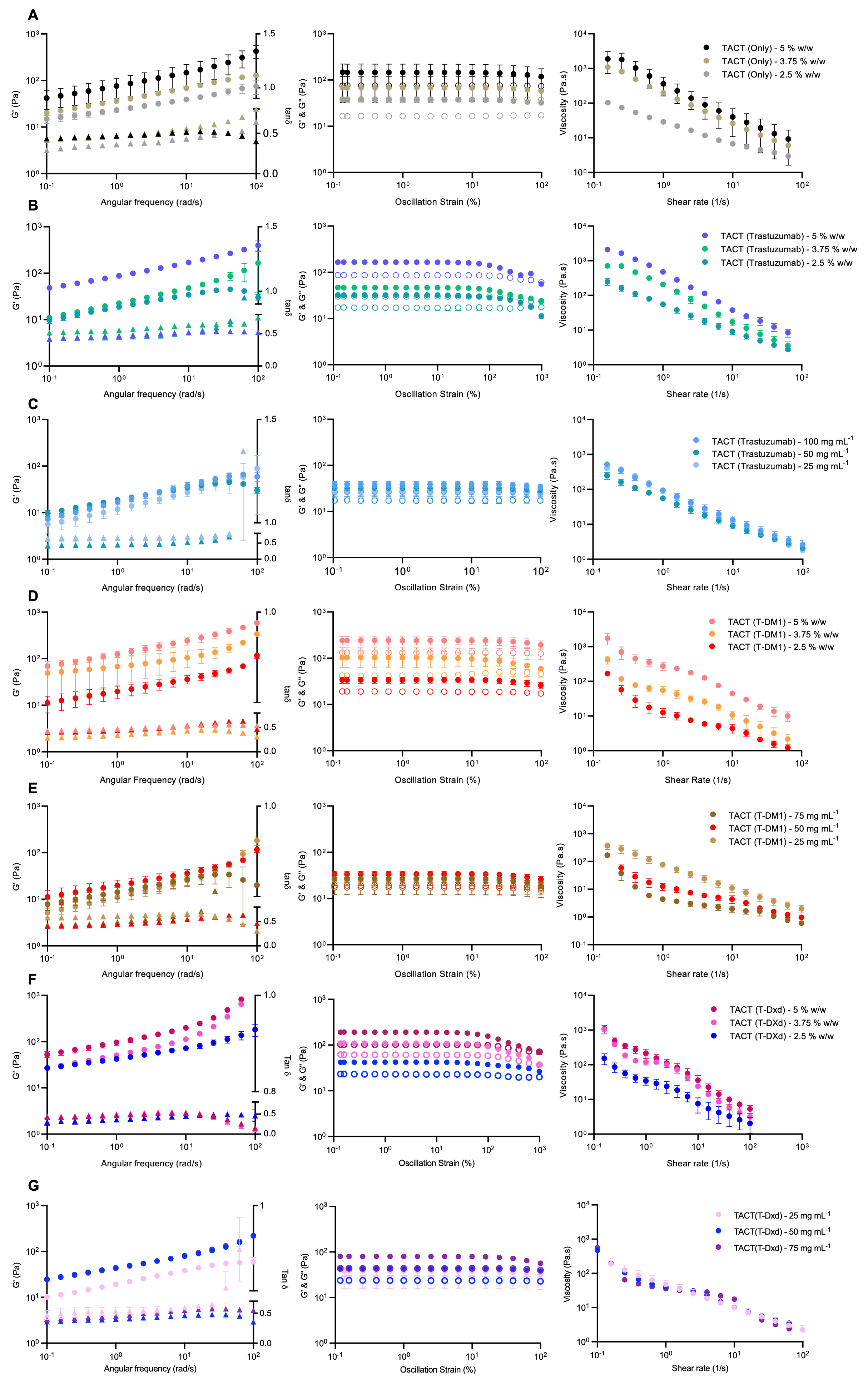

**Supplementary Figure 2. Extended rheological characterization of TACT formulations.**

Extended rheological characterization (at 25°C) based on TACT polymer concentration (% w/w) and antibody loading (mg mL^-1^). From left to right: frequency sweeps (filled circles represent the storage modulus and filled triangles represent tan delta), strain sweeps (filled and empty circles represent respectively the storage and loss modulus) and flow curves for **A.** TACT (Only), **B.** TACT(trastuzumab) at 50 mg mL-1 antibody, **C.** TACT(trastuzumab) at 2.5 % w/w polymer, **D.** TACT(T-DM1) at 50 mg mL^-1^ ADC, **E.** TACT(T-DM1) at 2.5 % w/w polymer, **F.** TACT(T-DXd) at 50 mg mL^-1^ ADC, **G.** TACT(T-DXd) at 2.5 % w/w polymer.

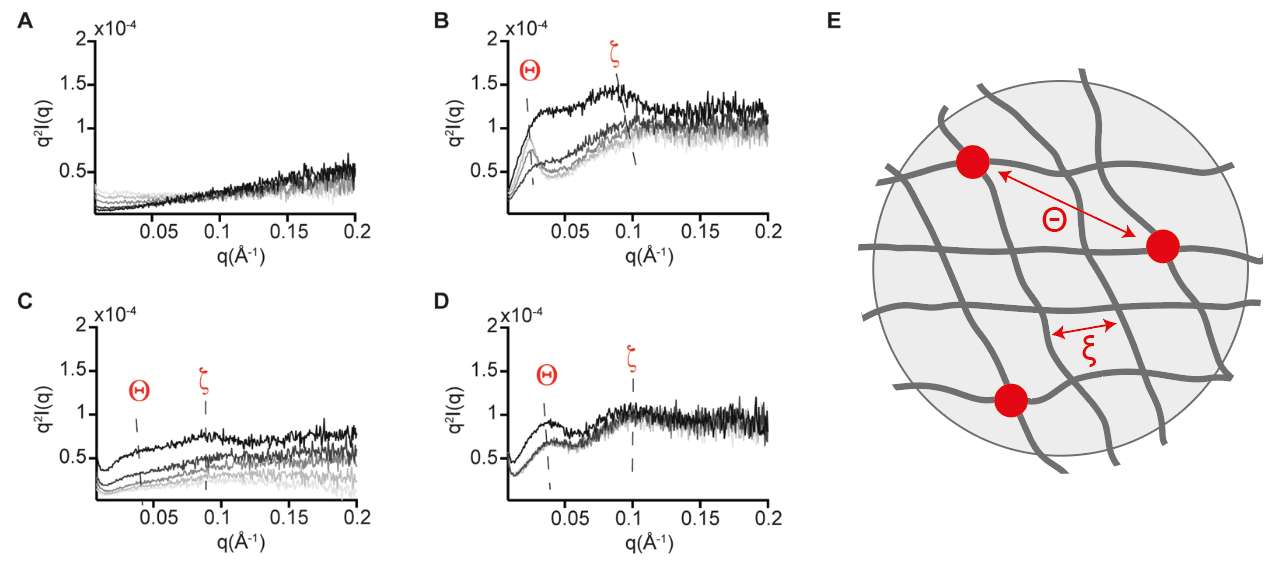

**Supplementary Figure 3.** Kratky plots (q^2^I(q) as a function of the wave vector q) for the 4 different gels. **A.** Kratky plot for the TACT alone. No specific correlation lengths are apparent during the 17 first hours. **B.** Kratky plot for TACT(trastuzumab). Two correlation lengths appear around 0.02A^-1^ and 0.09A^-1^ (dashed lines). **C.** Kratky plot for TACT(T-DM1). Two correlation length appear around 0.02A^-1^ and 0.09A^-1^ (dashed lines). **D.** Kratky plot for TACT(T-Dxd). Two correlation length appear around 0.03A^-1^ and 0.1A^-1^ (dashed lines). **E.** Scheme of the network formed when the 2-correlation length are present.

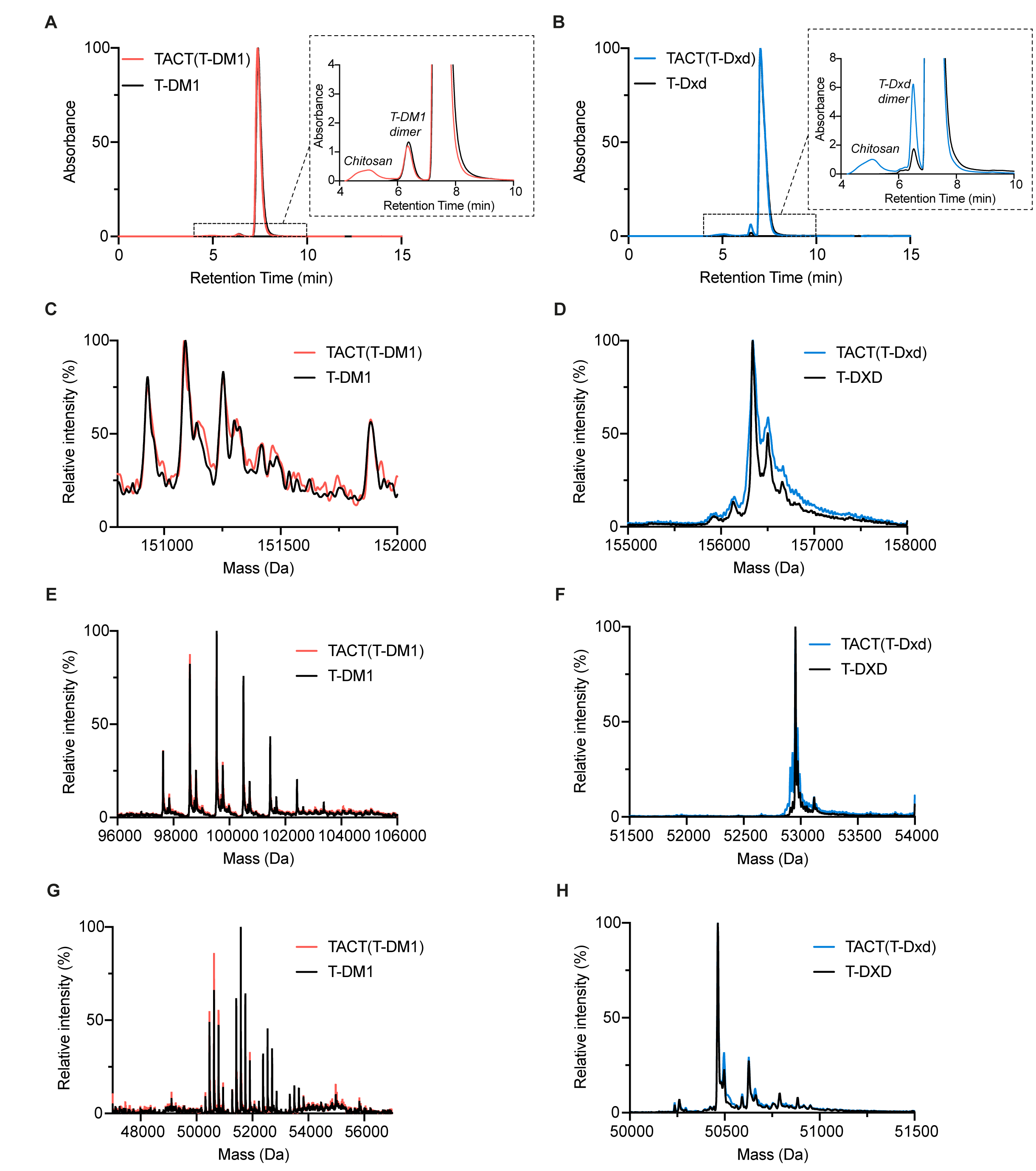

**Supplementary Figure 4.** Extended ADC integrity assessment after gel residence by SEC-nMS and middle-up MS for T-DM1 and T-DXd. SEC-UV chromatograms (280 nm) of **A.** T-DM1 and **B.** T-DXd. **C.** T-DM1 and **D.** T-DXd glycoforms distribution by intact SEC-nMs. **E.** T-DM1 F(ab’)_2_ fragment and **F.** T-DXd Fab fragment MS spectra by middle-up analysis. **G.** T-DM1 Fc sub-unit and **H.** T-DXd Fc sub-unit MS spectra by middle-up analysis. Fresh ADC (black) and TACT-released ADC (red for T-DM1 and blue for T-DXd).

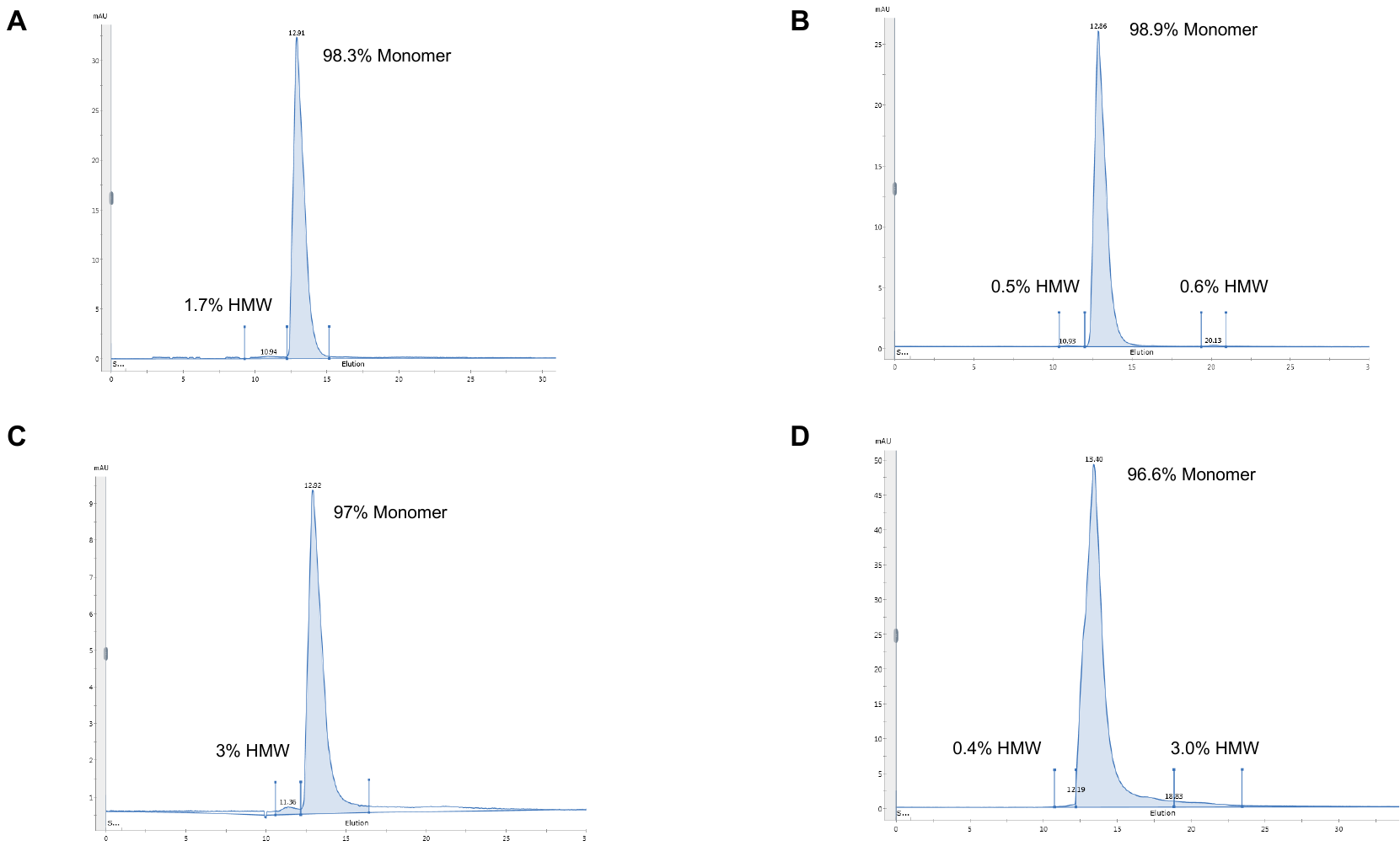

**Supplementary Figure 5.** Size-exclusion chromatography (SEC) chromatogram of **A.** trastuzumab, **B.** T-MMAE DAR=2, **C.** T-MMAE DAR=3.8 and **D.** T-MMAE DAR=8 (2 mg mL^-1^, 50 µL). Analyses were performed with UV detection at 280 nm. Peaks corresponding to high-molecular-weight (HMW) species, monomer, and low-molecular-weight (LMW) species are indicated.

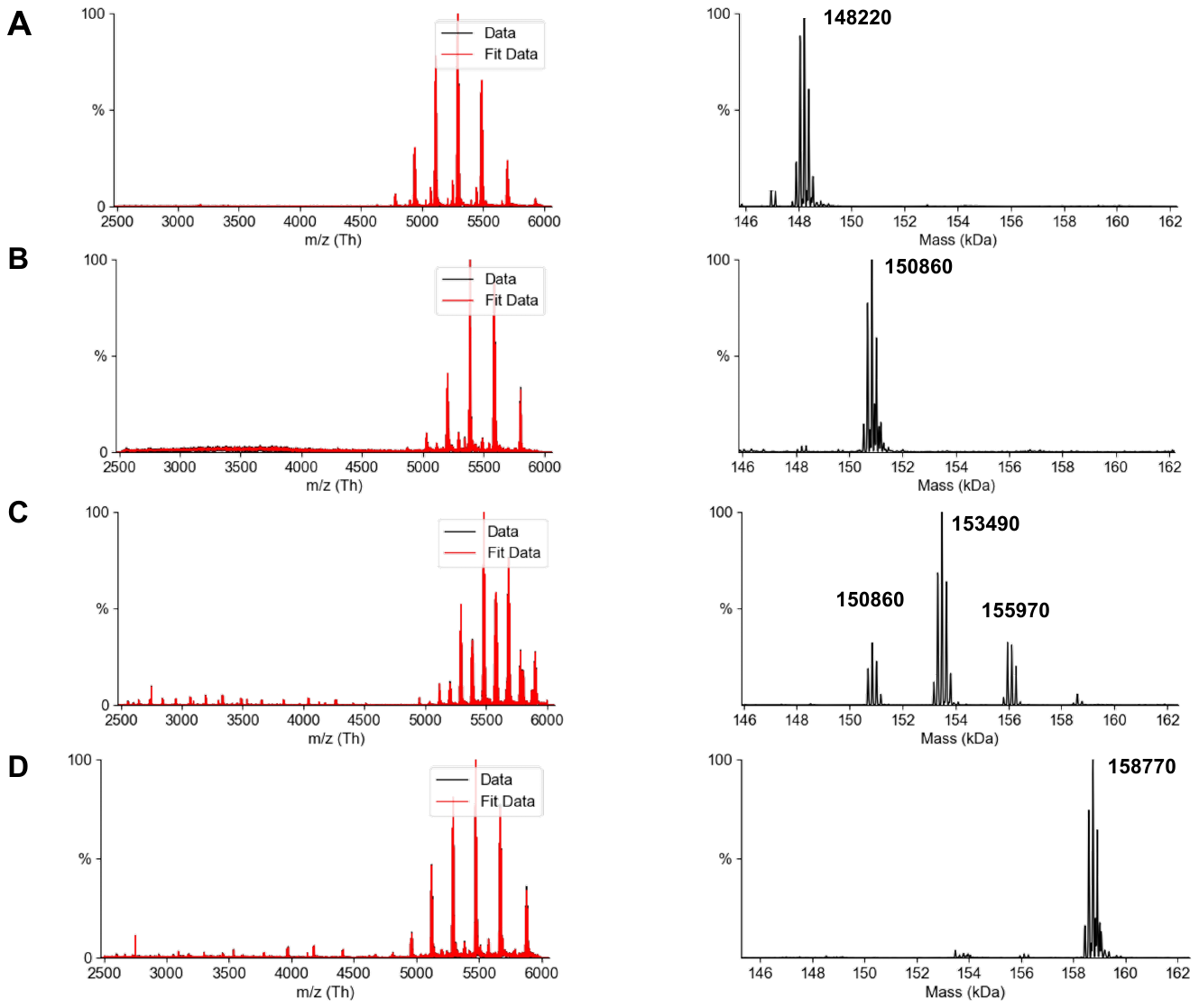

**Supplementary Figure 6**. DAR distribution of the unmodified trastuzumab and MMAE-conjugates by size exclusion chromatography (SEC) - native mass spectra analysis. Native mass spectra (left) and corresponding mass distributions (right) are shown for **A.** trastuzumab, **B.** T-MMAE DAR=2, **C.** T-MMAE avDAR=3.8 and **D.** T-MMAE DAR=8.

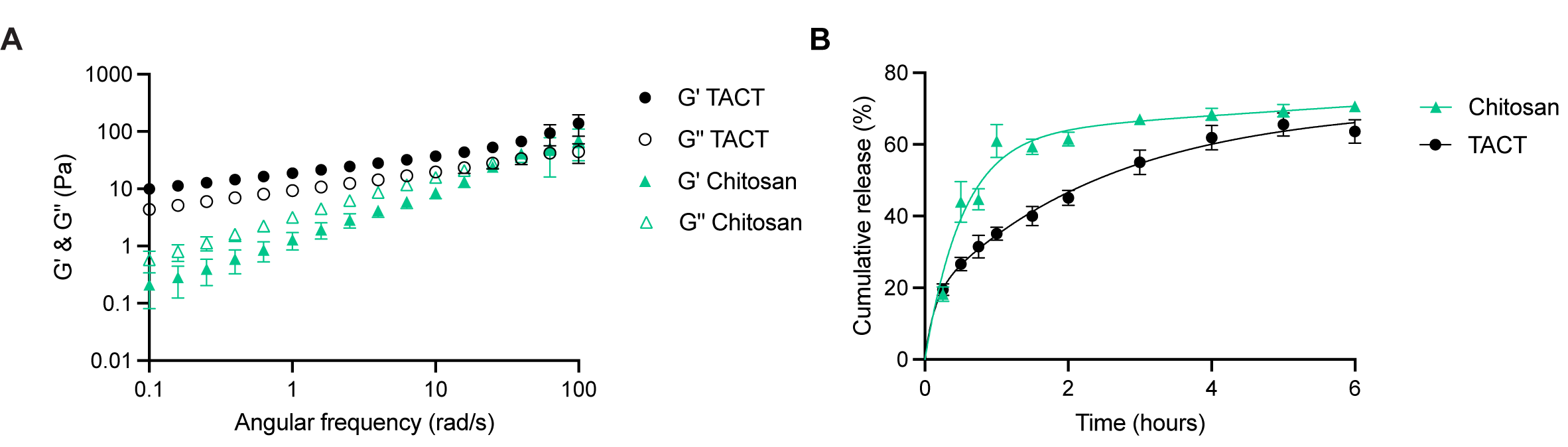

**Supplementary Figure 7.** **A.** Rheological comparison showing TACT (G’ > G’’) versus unmodified chitosan (G’ < G’’). **B.** Cumulative release profiles of TACT(trastuzumab) and chitosan-trastuzumab in SC simulated media (n = 3, mean ± SEM).

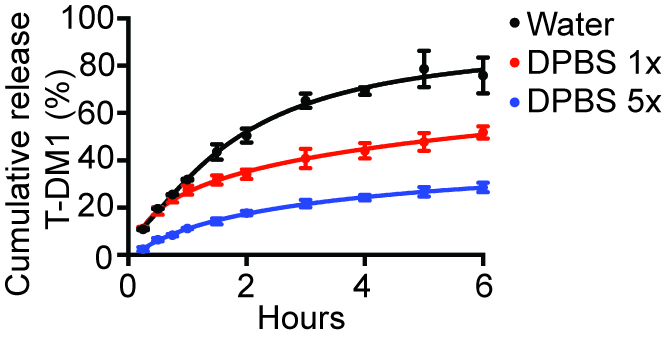

**Supplementary Figure 8.** Ionic-strength dependence of T-DM1 release from TACT. Cumulative release profiles in water, 1X DPBS and 5X DPBS (n = 2-3, mean ± SEM, 25 °C).

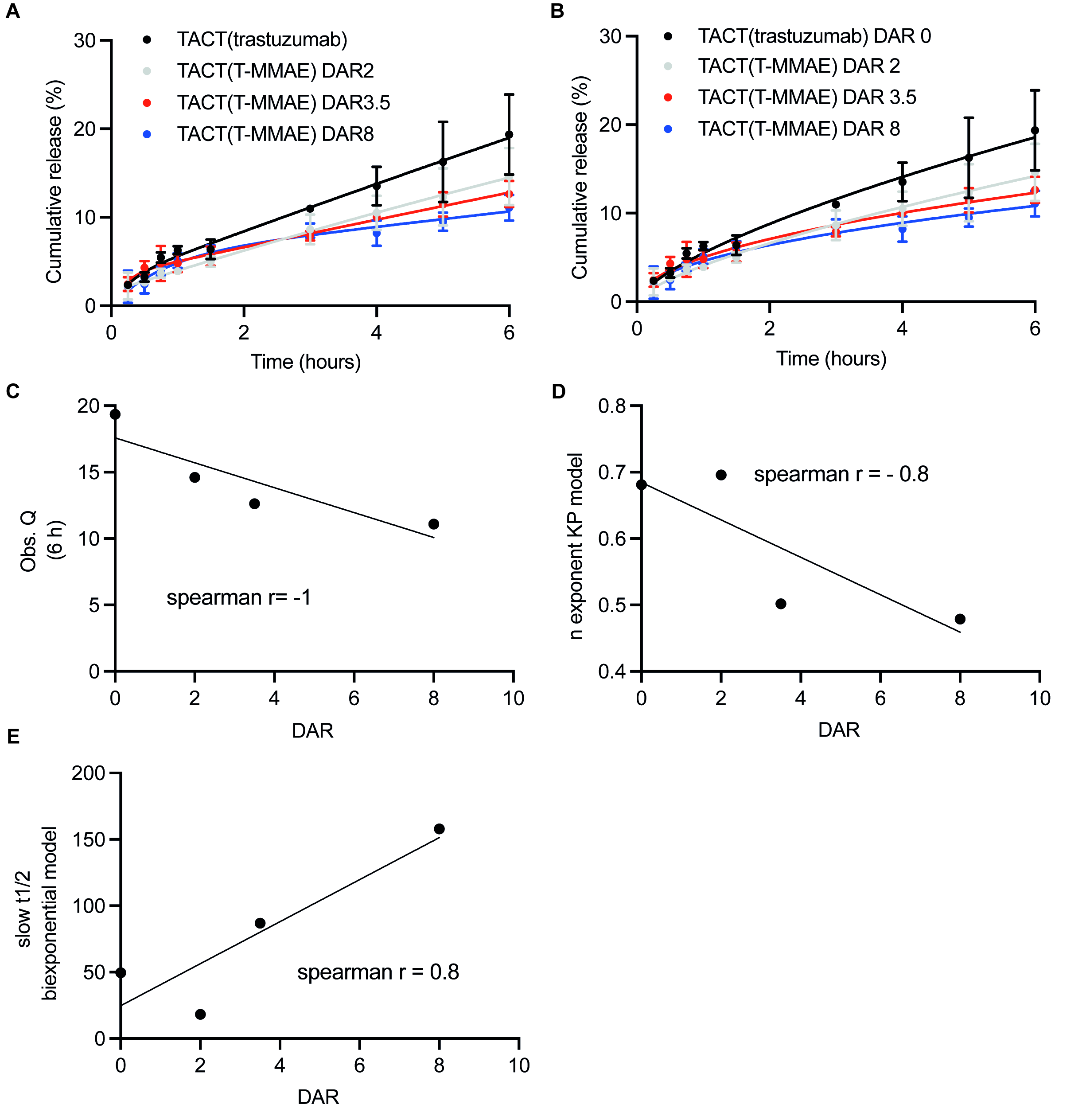
**Supplementary Figure 9.** Kinetic modeling (Korsmeyer-Peppas and biexponential) of the trastuzumab-MMAE DAR series (DAR 0, 2, 3.5, 8). **A.** Cumulative release profiles with bi-exponential fit. **B.** Cumulative release profiles with Korsmeyer-Peppas fit. **C.** Cumulative release (Obs. Q) at 6h monotonically decreases with DAR (spearman coefficient = 1). **D.** Exponent n (Korsmeyer-Peppas model) versus DAR, with a shift toward Fickian diffusion at higher DAR (spearman coefficient r = -0.8). **E.** Slow half-life (t_1/2_, bi-exponential model) increases with DAR except DAR 2 (spearman coefficient r= 0.8).

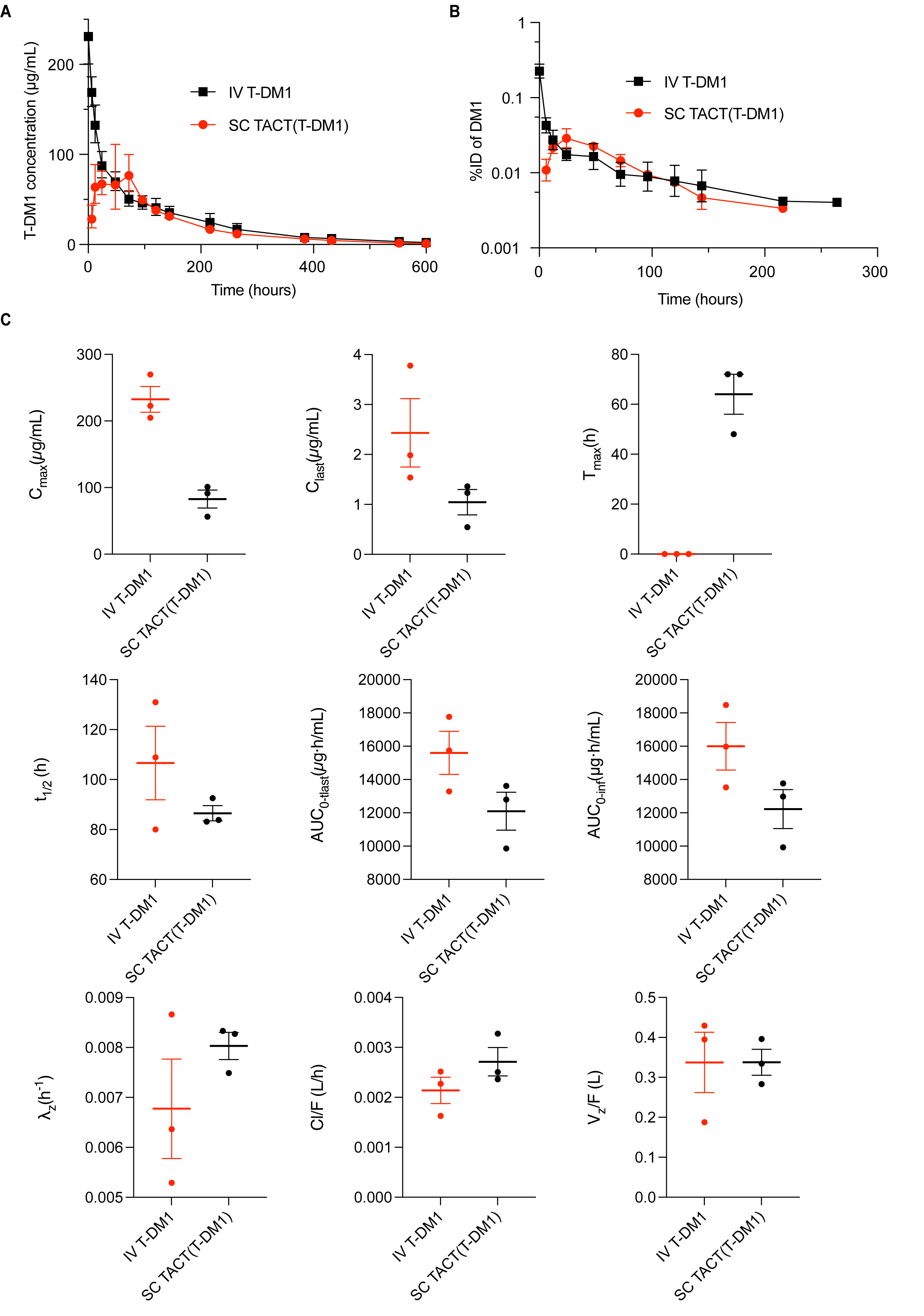
**Supplementary Figure 10.** NHP pharmacokinetic profiles and systemic exposure. **A.** Mean plasma concentration-time profiles for intact T-DM1 after IV and SC TACT administration (n = 3 per group). **B.** DM1 catabolite profiles after IV and SC TACT administration (n = 3 per group), expressed as percentage of injected dose (ID). **C.** Noncompartmental analysis parameter summary between IV T-DM1 and SC TACT(T-DM1).

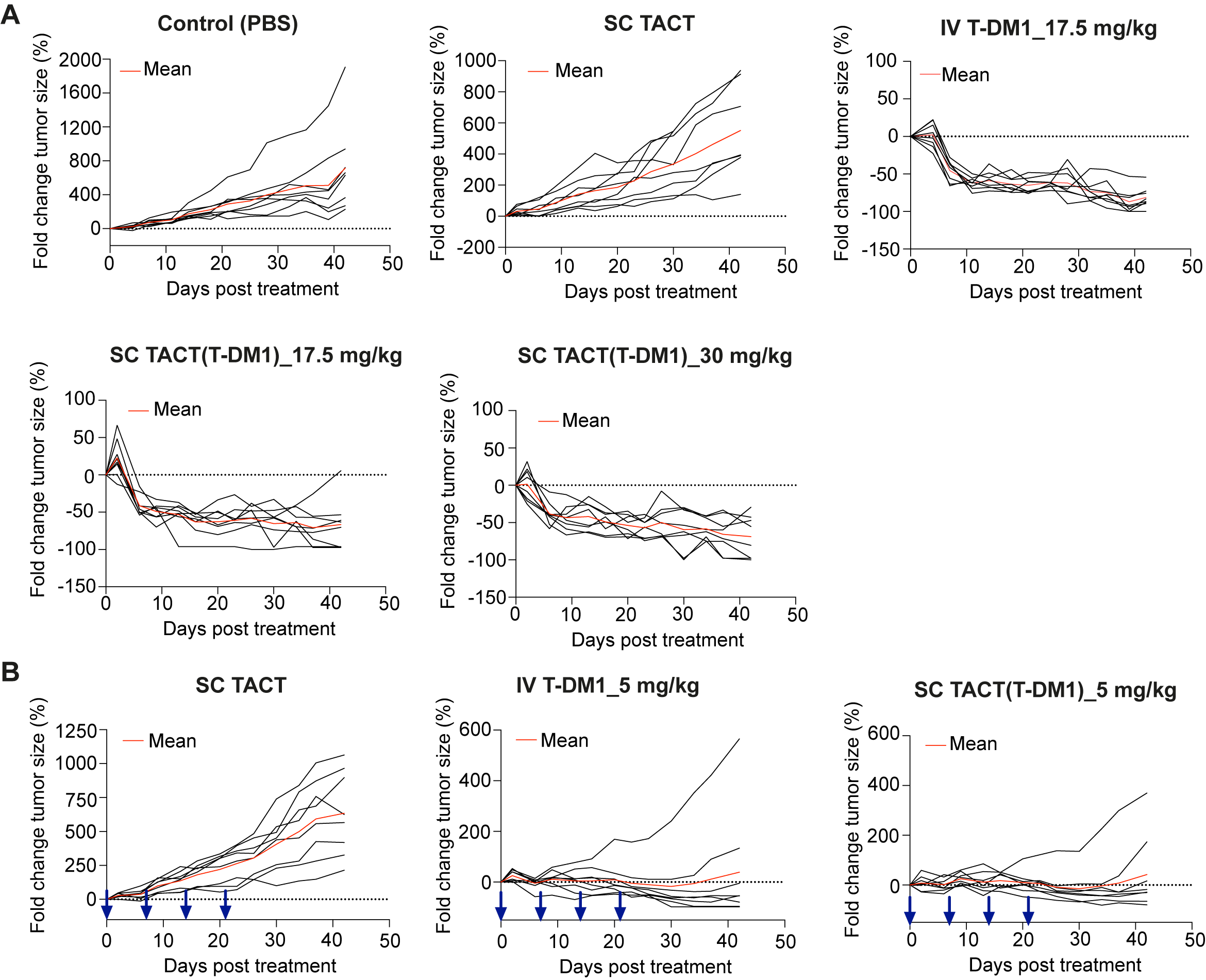

**Supplementary Figure 11.** Individual tumor growth curves (spider plots) for **A.** T-DM1 single-dose and **B.** T-DM1 multi-dose (Q7D×4). Each line represents one animal and mean value is represented in red.

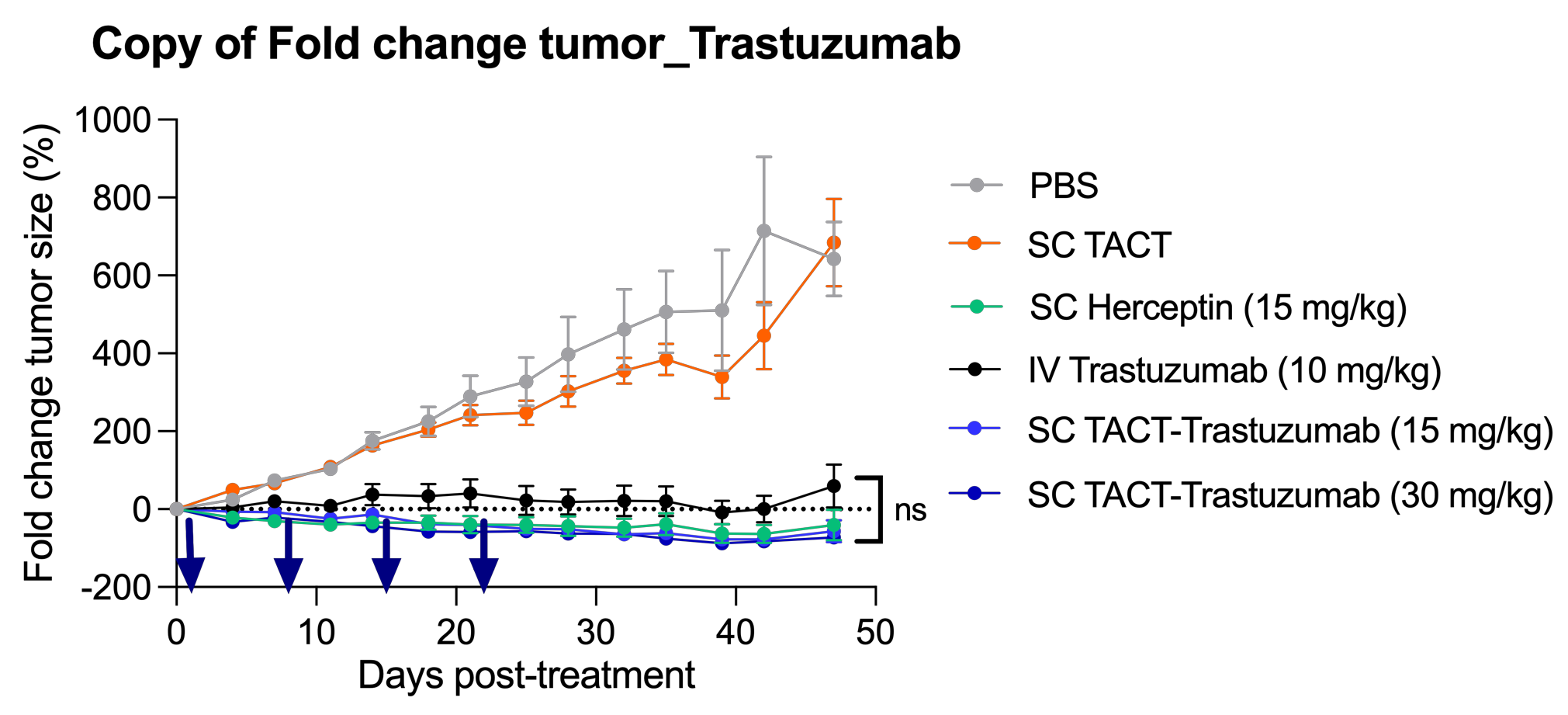

**Supplementary Figure 12.** Antitumor efficacy of trastuzumab formulations. Q7D×4 treatment in BT-474 xenograft mice comparing IV Trazimera (10 mg kg^-1^), SC Herceptin (15 mg kg^-1^) and SC TACT(trastuzumab) (15 and 30 mg kg^-1^; n = 8 per group). Data are mean ± SEM and arrows represent treatment injections.

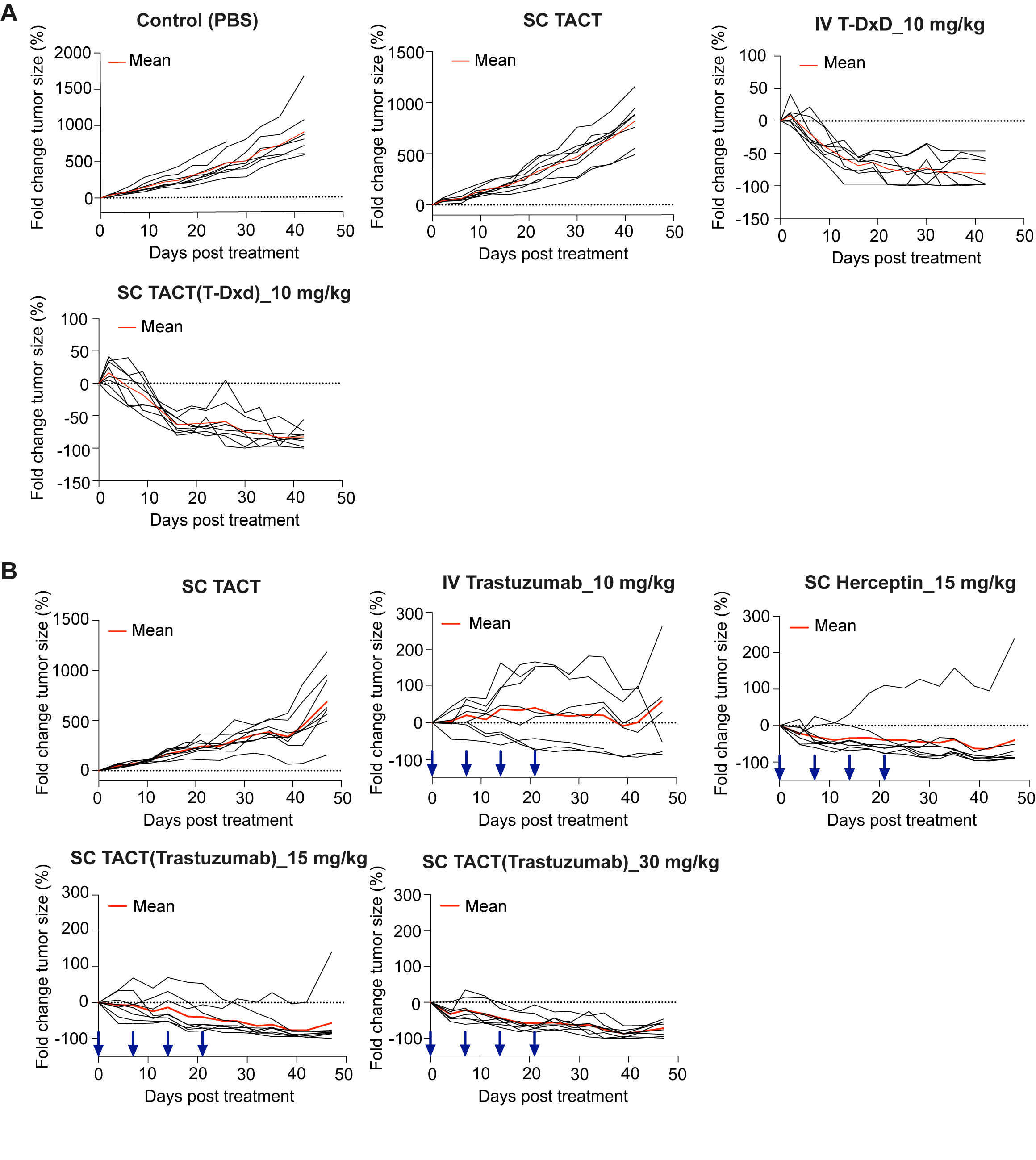

**Supplementary Figure 13.** Individual tumor growth curves (spider plots) for **A.** T-DXd single-dose and **B.** trastuzumab multi-dose (Q7D×4) efficacy studies. Each line represents one animal and mean value is represented in red.

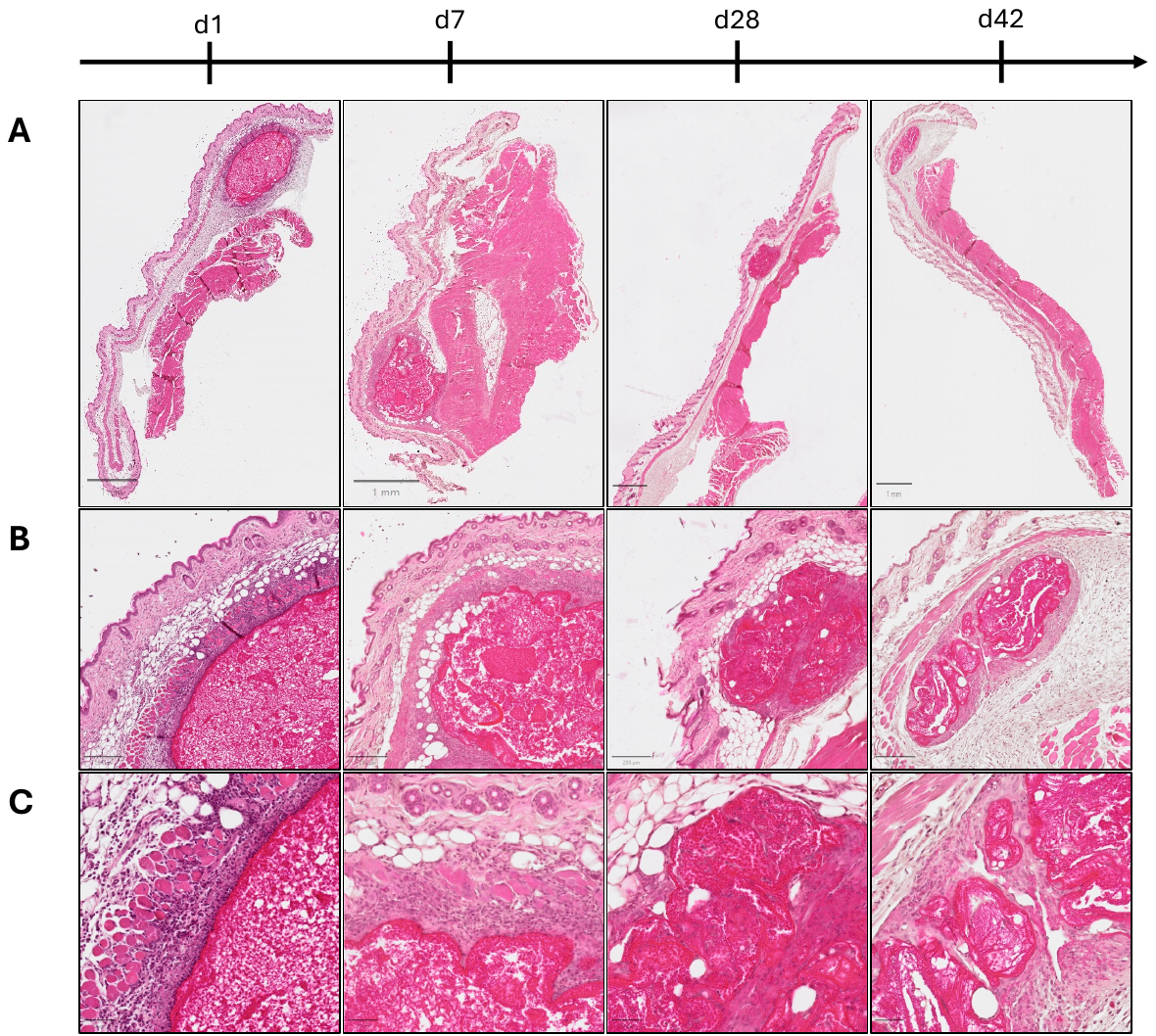

**Supplementary Figure 14. Histological assessment of depot integration and host-cell ingress over time.** H&E staining of TACT(T-DM1) injection sites at days 1, 7, 28 and 42 post injection. Three magnification levels are shown for each time point: **A**. Overview of the injection site (scale bar, 1 mm), **B**. Intermediate magnification showing skin layers and cellular infiltration (scale bar, 200 µm), **C**. High-magnification view of the shell-gel interface (scale bar: 50 µm). This figure shows progressive immune-cell infiltration at the depot periphery without disruption of surrounding tissue architecture.

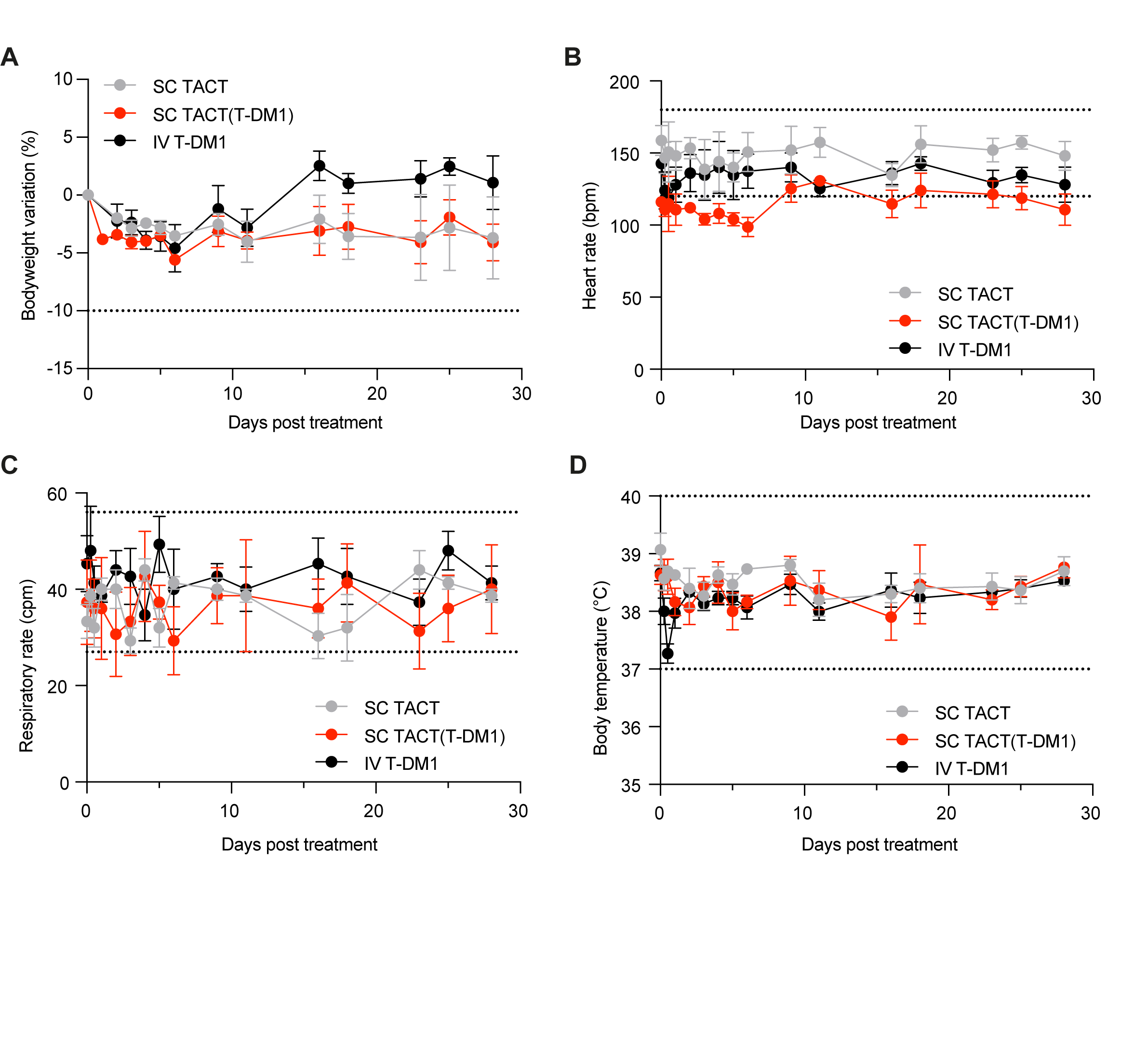

**Supplementary Figure 15.** NHP vital signs. **A.** Body weight, **B.** heart rate, **C.** respiratory rate and **D.** body temperature for all three groups (TACT hydrogel alone, TACT(T-DM1), T-DM1 IV; n = 3 per group). Data are represented as mean ± SEM.

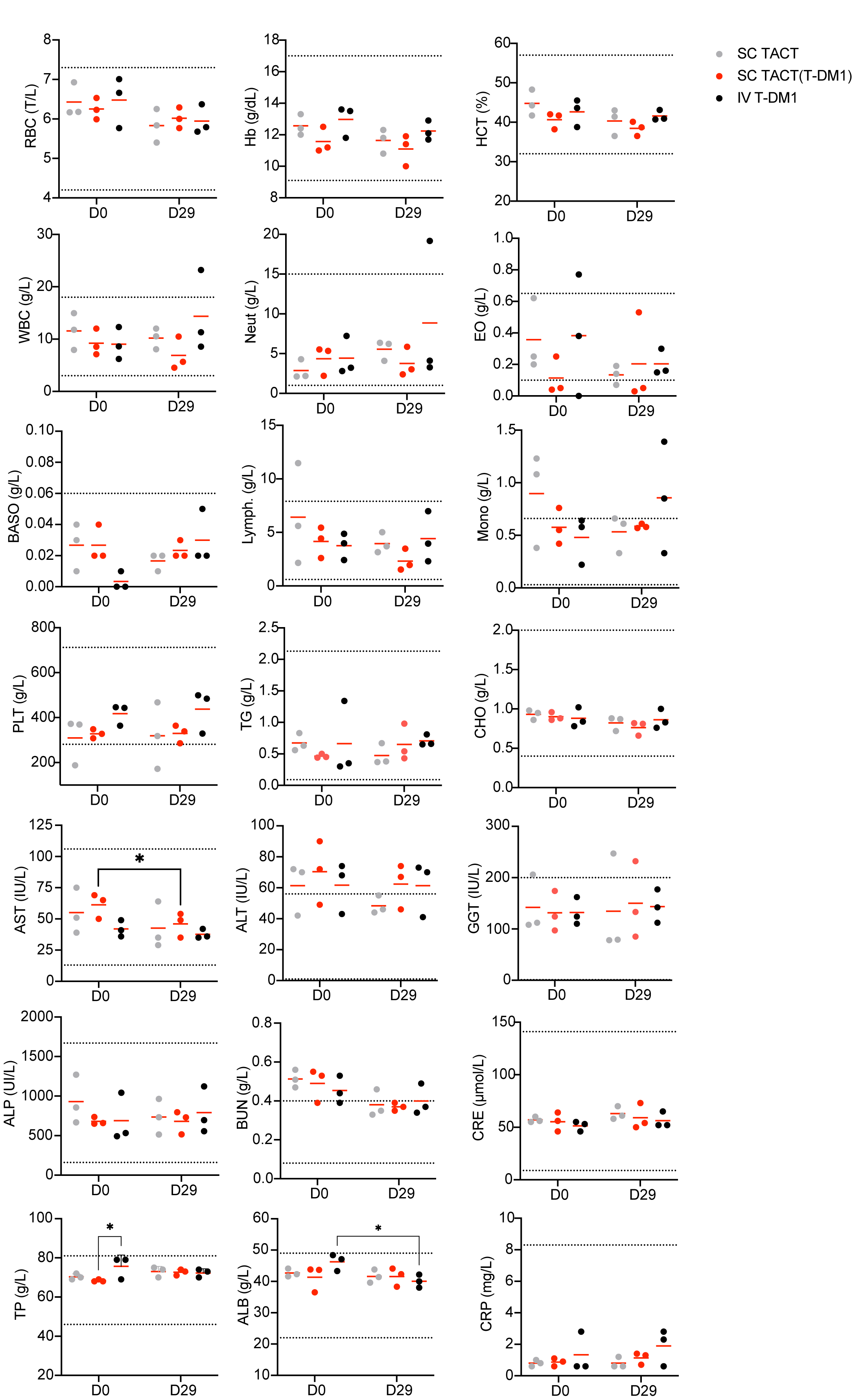

**Supplementary Figure 16.** NHP hematological and biochemical parameters. Box plots of red blood cells (RBC), hemoglobin (Hb), hematocrit (HCT), white blood cells (WBC), neutrophils (Neut), eosinophils (EO), basophils (BASO), lymphocytes (Lymph.), monocytes (Mono), platelet counts (PLT), triglycerides (TG), cholesterol (CHO), hepatic transaminases (AST and ALT), gamma glutamyl transferase (GGT), alkaline phosphatase (ALP), blood urea nitrogen (BUN), creatinine (CRE), total proteins (TP), albumin (ALB) and C-reactive protein (CRP) for all three groups (TACT hydrogel alone, TACT(T-DM1), T-DM1 IV; n = 3 per group) at baseline and day 26. Individual data points shown, with mean value. Dotted lines represent the expected physiological range.

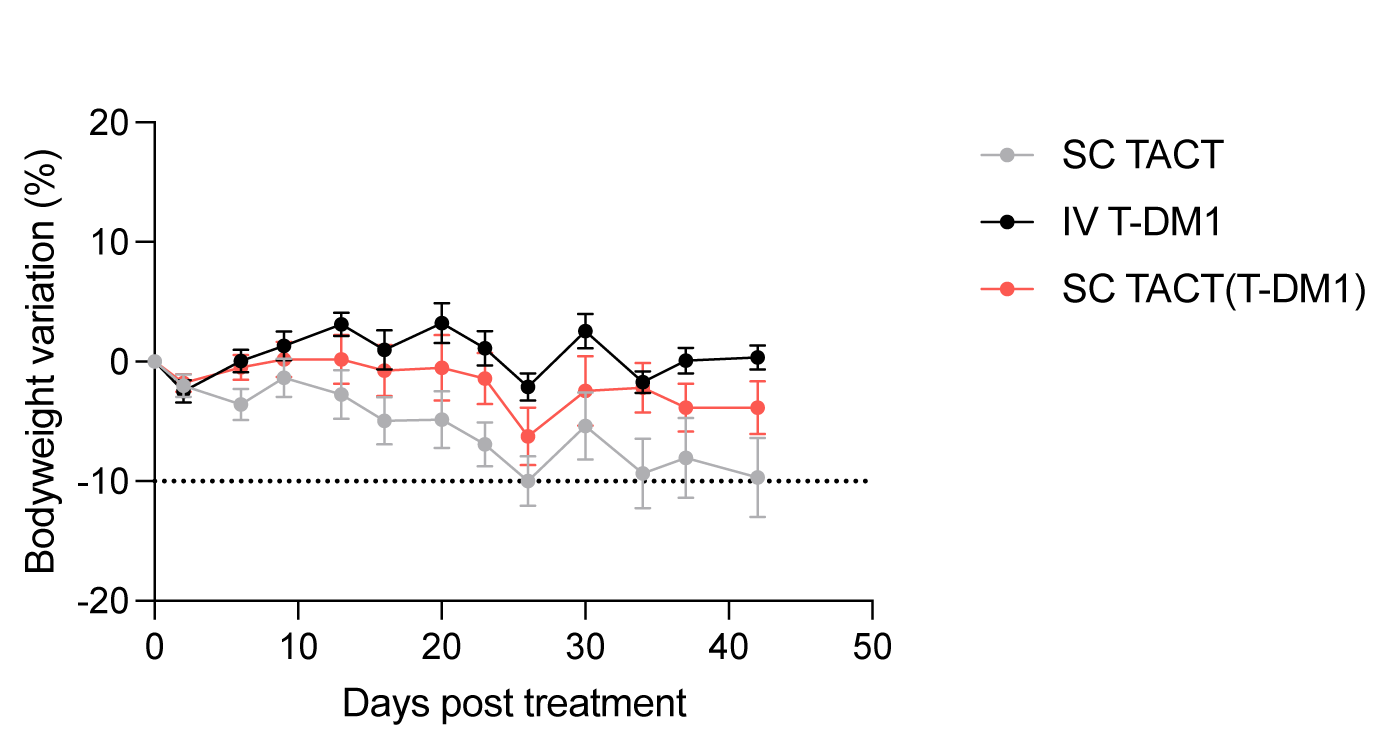

**Supplementary Figure 17.** Local tolerance after repeated SC dosing. Body weight monitoring in xenograft tumor mice model (BT-474) receiving Q7D×4 SC TACT hydrogel, IV T-DM1 or SC TACT(T-DM1) (5 mg kg^-1^ of T-DM1 per dose).

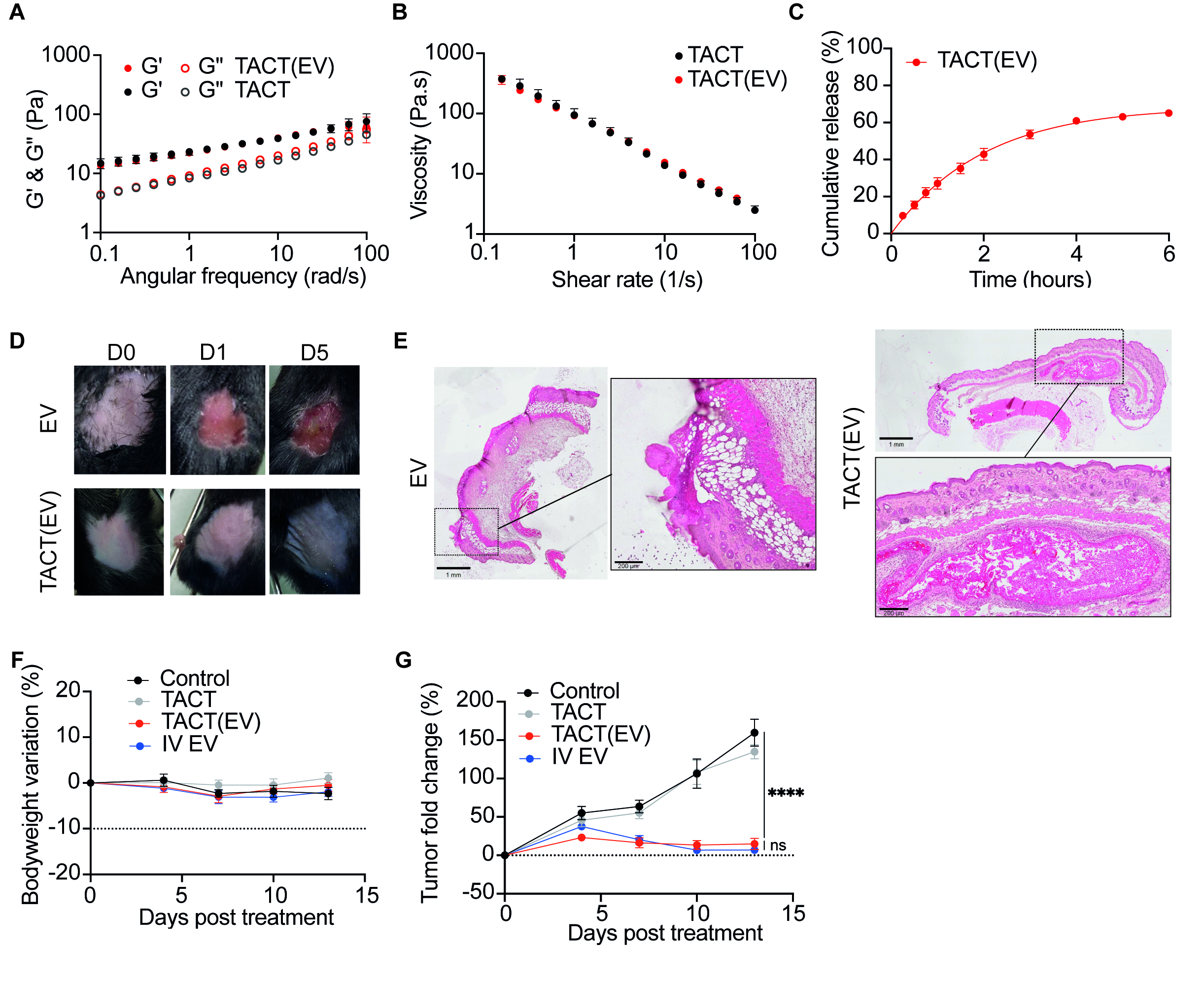
**Supplementary Figure 18.** Demonstration of TACT hydrogel versatility on a second irritant ADC (enfortumab vedotin, EV). **A.** Frequency sweep of TACT (black) and TACT(EV) (red) at 2.5% w/w TACT and 50 mg mL^-1^ EV. **B.** Flow curve of TACT (black) and TACT(EV) (red) at 2.5% w/w TACT and 50 mg mL^-1^ EV. **C.** Cumulative release profiles of TACT(EV) in simulated SC media. **D.** Macroscopic photographs of SC injection sites in C57BL/6 mice over 5 days for EV alone and TACT(EV). **E.** Histopathology (H&E staining) of injection sites at day 5 for EV alone and TACT(EV). **F.** Body weight monitoring of HT-1376 xenograft-bearing mice over the treatment period. **G.** Tumor growth in HT-1376 xenograft-bearing mice treated with a single dose of SC TACT(EV) or IV EV (10 mg kg^-1^; n = 8 per group). Data are mean ± SEM. ****P < 0.0001 versus control; ns, not significant between SC and IV treated groups.

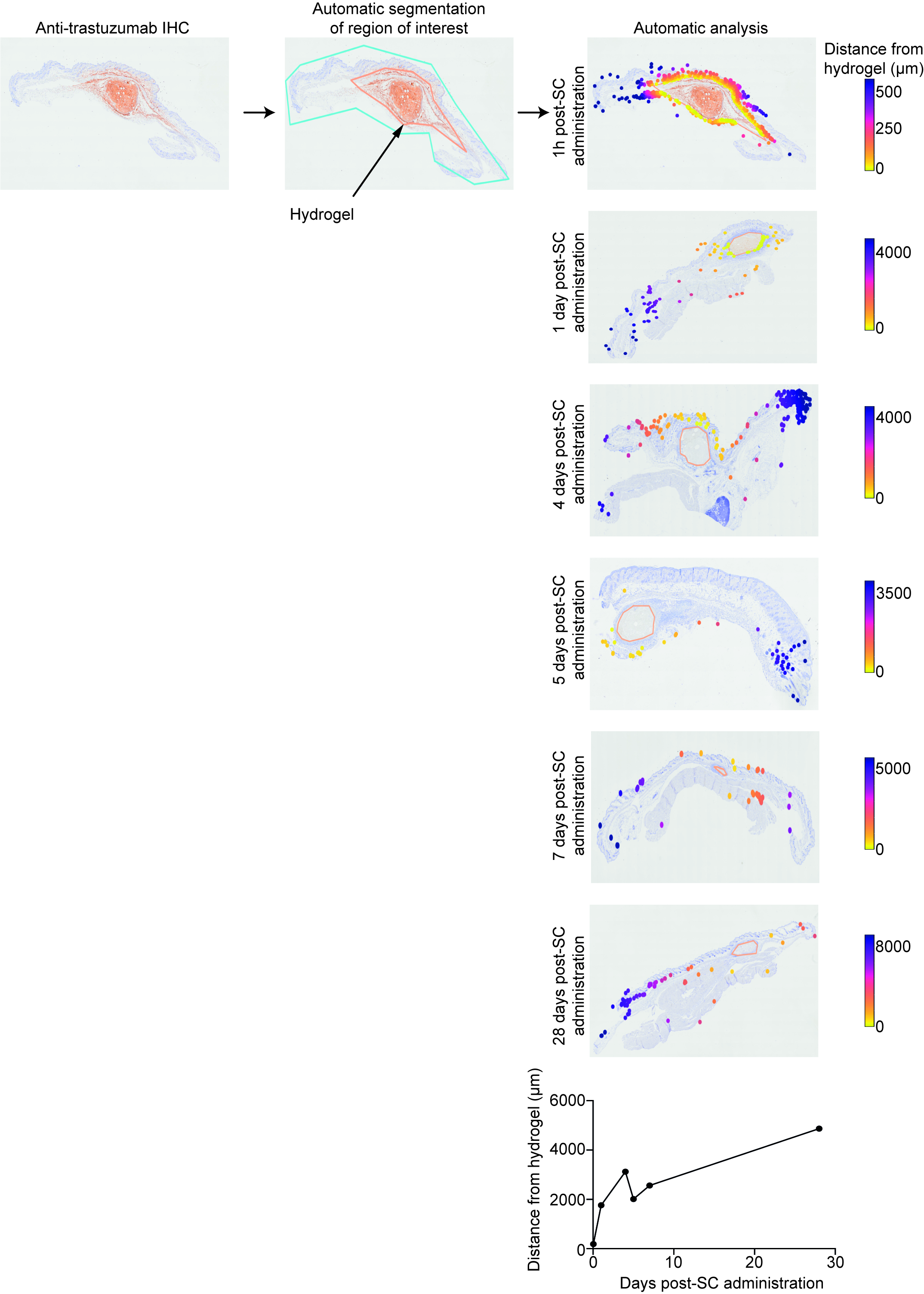
**Supplementary Figure 19.** Extended anti-trastuzumab immunohistochemistry of SC injection sites following SC TACT(T-DM1) delivery. Representative whole-slide anti-trastuzumab staining over the acute, intermediate and resolved phases of depot evolution for TACT(T-DM1) to characterize the distance from the implantation site.

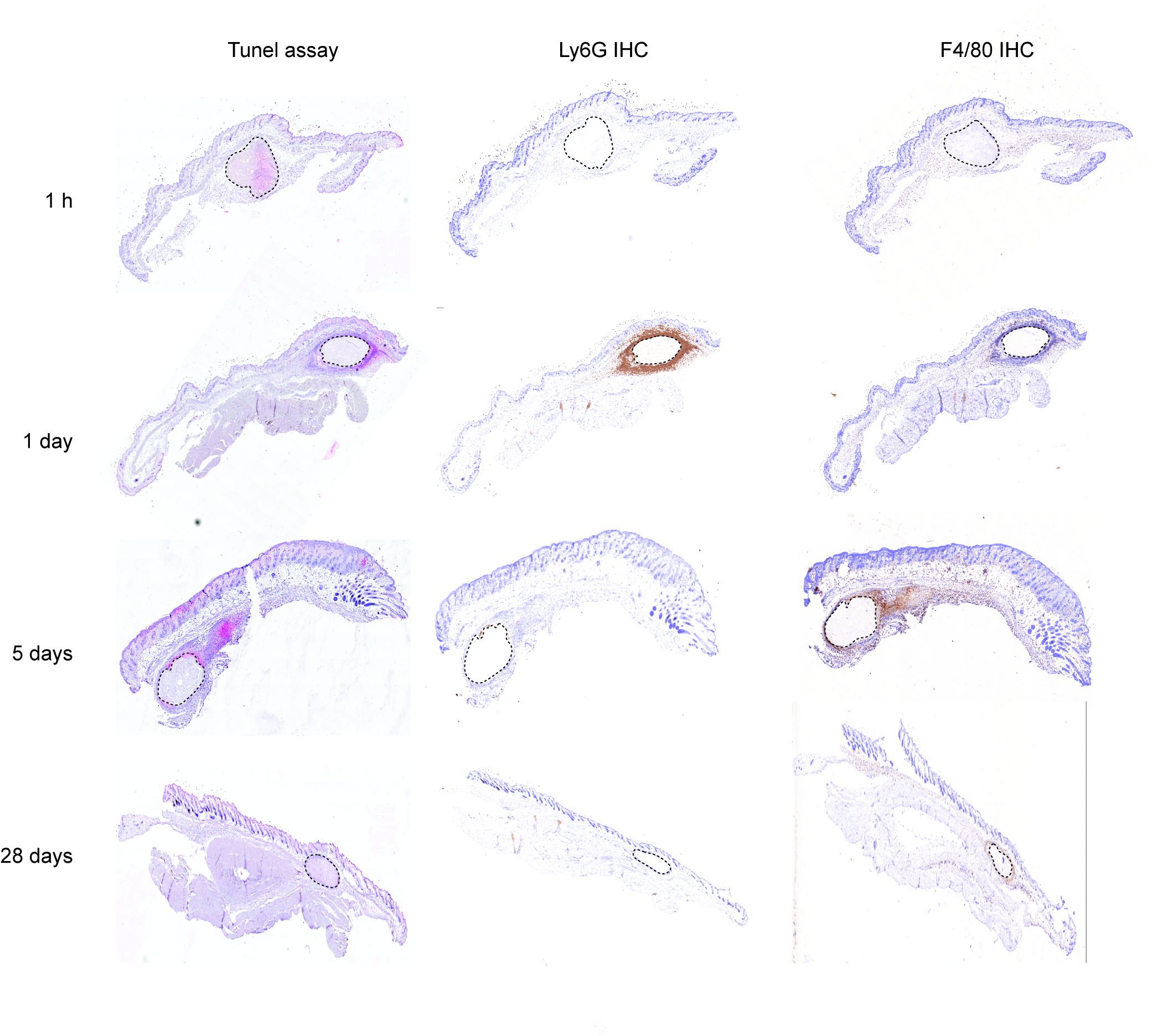

**Supplementary Figure 20.** Longitudinal immunohistochemistry (IHC) staining of TACT(T-DM1) in healthy C57BL/6 mice.

**Section 4. Supplementary Tables**

**Supplementary Table 1. Summary of TACT formulation metrics (polymer concentration, G', extrusion force).**

| **Formulation** | **Polymer concentration (% w/w)** | **Antibody concentration (mg mL^-1^)** | **G’ (Pa) at 10 rad s^-1^** | **Tan𝛿**  **at 10 rad s^-1^** | **Extrusion force (N) at 1 mL min^-1^** |
| --- | --- | --- | --- | --- | --- |
| **TACT** | 2.5 | 0 | 39 ± 4 | 0.43 ± 0.03 | 6.9 ± 0.4 |
|  | 3.75 | 0 | 69 ± 5 | 0.56 ± 0.02 | ND |
|  | 5.0 | 0 | 148 ± 74 | 0.52 ± 0.03 | ND |
| **TACT(T-DM1)** | 2.5 | 25 | 25 ± 6 | 0.66 ± 0.05 | ND |
|  |  | 50 | 36 ± 7 | 0.55 ± 0.10 | 6.0 ± 0.4 |
|  |  | 75 | 27 ± 8 | 0.66 ± 0.01 | ND |
|  | 3.75 | 50 | 105 ± 41 | 0.44 ± 0.18 | ND |
|  | 5.0 | 50 | 244 ± 48 | 0.54 ± 0.04 | ND |
| **TACT(T-DXd)** | 2.5 | 25 | 38 ± 8 | 0.59 ± 0.06 | ND |
|  |  | 50 | 79(σ n.d.) | 0.46(σ n.d.) | 9.9 ± 0.1 |
|  |  | 75 | 81(σ n.d.) | 0.55(σ n.d.) | ND |
|  | 3.75 | 50 | 113 ± 3 | 0.54 ± 0.01 | ND |
|  | 5.0 | 50 | 199 ± 22 | 0.53 ± 0.01 | ND |
| **TACT(Trastuzumab)** | 2.5 | 25 | 27 ± 5 | 0.68 ± 0.10 | ND |
|  |  | 50 | 34 ± 0 | 0.52 ± 0.06 | 7.0 ± 0.0 |
|  |  | 75 | 33 ± 2 | 0.51 ± 0.05 | ND |
|  |  | 100 | 38 ± 6 | 0.54 ± 0.08 | ND |
|  | 3.75 | 50 | 48 ± 1 | 0.63 ± 0.01 | ND |
|  | 5.0 | 50 | 170 ± 1 | 0.52 ± 0.01 | ND |

Extrusion forces were measured in air, using 1mL syringe with a 25G needle (25G, 16 mm).

**Supplementary Table 2. Cryo-SEM pore-size summary statistics across conditions (at 40k magnification).**

| **Condition** | **N images** | **N pores** | **Median (nm)** | **Mean ± SD** | **Pore density (pores/mm^2^)** | **Circularity mean ± SD** |
| --- | --- | --- | --- | --- | --- | --- |
| **TACT,** *water* | 5 | 2704 | 69.1 | 72.0 ± 40.4 | 78.08 ± 5.80 | 0.650 ± 0.013 |
| **TACT,** *PBS* | 5 | 3112 | 55.6 | 62.8 ± 38.6 | 89.85 ± 4.37 | 0.675 ± 0.013 |
| **TACT(T-DM1),** *PBS* | 4 | 2037 | 55.8 | 65.7 ± 42.2 | 70.48 ± 2.30 | 0.657 ± 0.016 |
| **TACT(T-DXd),** *PBS* | 3 | 1761 | 56.6 | 63.4 ± 37.9 | 84.70 ± 3.62 | 0.661 ± 0.001 |

| **Condition** | **N pores** | **Median (nm)** | **Mean (nm)** | **SD (nm)** | **Density (pores/µm²)** | **Circularity** |
| --- | --- | --- | --- | --- | --- | --- |
| **TACT,** *water* | 501 | 75.0 | 74.8 | 40.7 | 72.29 | 0.641 |
|  | 514 | 70.9 | 75.4 | 43.2 | 74.17 | 0.656 |
|  | 555 | 68.9 | 69.8 | 37.7 | 80.08 | 0.632 |
|  | 532 | 66.6 | 72.1 | 42.4 | 76.84 | 0.663 |
|  | 602 | 64.1 | 68.3 | 38.0 | 87.02 | 0.659 |
| **TACT,** *PBS* | 655 | 57.5 | 61.2 | 36.8 | 94.43 | 0.675 |
|  | 635 | 52.2 | 60.8 | 38.2 | 91.54 | 0.682 |
|  | 589 | 63.1 | 66.9 | 38.8 | 84.99 | 0.660 |
|  | 592 | 52.5 | 64.5 | 41.1 | 85.42 | 0.665 |
|  | 641 | 52.8 | 60.9 | 38.2 | 92.88 | 0.692 |
| **TACT(T-DM1),** *PBS* | 508 | 53.4 | 63.8 | 41.6 | 73.30 | 0.664 |
|  | 532 | 58.8 | 66.5 | 40.7 | 76.76 | 0.652 |
|  | 501 | 60.7 | 69.6 | 43.8 | 72.29 | 0.638 |
|  | 496 | 50.4 | 62.7 | 42.7 | 71.57 | 0.675 |
| **TACT(T-DXd),** *PBS* | 616 | 57.9 | 63.0 | 36.5 | 88.88 | 0.661 |
|  | 573 | 55.8 | 63.7 | 39.1 | 82.68 | 0.663 |
|  | 572 | 56.1 | 63.5 | 38.2 | 82.53 | 0.660 |

**Supplementary Table 3. T-DM1 aggregation state by SEC-UV (280 nm) post release from TACT hydrogel, compared to T-DM1 reference**.

| **Conditions** | **%HMWS** | **%Monomer** | **%LMWS** |
| --- | --- | --- | --- |
| T-DM1 | 1.5 ± 0.1 | 98.5 ± 0.1 | ND |
| TACT(T-DM1) | 2.4 ± 0.1 | 97.6 ± 0.1 | ND |

HMWS: High molecular weight species; LMWS: Low molecular weight species; ND: Not quantifiable.

**Supplementary Table 4. T-DXd aggregation state by SEC-UV (280 nm) post release from TACT hydrogel, compared to T-DXd reference**.

| **Conditions** | **%HMWS** | **%Monomer** | **%LMWS** |
| --- | --- | --- | --- |
| T-DXd | 0.8 ± 0.1 | 99.2 ± 0.1 | ND |
| TACT(T-DXd) | 5.9 ± 0.4 | 94.1 ± 0.4 | ND |

HMWS: High molecular weight species; LMWS: Low molecular weight species; ND: Not quantifiable.

A fraction of the observed increase in HMWS content of TACT(T-DXd) can be attributed to the release medium used to extract T-DXd from the gel matrix. PBS was employed rather than the clinically recommended diluent for intravenous administration (5% glucose solution), as T-DXd exhibits a known propensity to aggregate in saline-based matrices. Despite this limitation, PBS was deliberately selected to more closely reflect physiological conditions.

**Supplementary Table 5. Distribution of T-DM1 glycoforms by intact SEC-MS post release from TACT hydrogel, compared to T-DM1 reference**.

| **Conditions** | **G0F/G0F (%)** | **G0F/G1F (%)** | **G1F/G1F (%)** |
| --- | --- | --- | --- |
| T-DM1 | 27.9 ± 2.0 | 39.5 ± 1.8 | 32.5 ± 2.1 |
| TACT(T-DM1) | 28.7 ± 1.3 | 39.5 ± 1.6 | 31.9 ± 0.7 |

**Supplementary Table 6. Distribution of T-DXd glycoforms by intact SEC-MS post release from TACT hydrogel, compared to T-DXd reference**.

| **Conditions** | **Unassigned (%)** | **A1G0F/A2G0F (%)** | **G0F/G0F (%)** | **G0F/G1F (%)** | **G1F/G1F (%)** |
| --- | --- | --- | --- | --- | --- |
| T-DXd | 3.0 ± 0.2 | 7 ± 0.2 | 51.9 ± 0.3 | 25.5 ± 0.5 | 12.6 ± 0.5 |
| TACT(T-DXd) | 3.5 ± 0.1 | 7.5 ± 0.1 | 47.3 ± 1.5 | 26.9 ± 1.0 | 14.7 ± 0.6 |

**Supplementary Table 7. Distribution of T-DM1 DAR species by intact SEC-MS post release from TACT hydrogel, compared to T-DM1 reference**.

| **Conditions** | **D0 (%)** | **D1 (%)** | **D2 (%)** | **D3 (%)** | **D4 (%)** | **D5 (%)** | **D6 (%)** | **D7 (%)** | **D8 (%)** | **avDAR** |
| --- | --- | --- | --- | --- | --- | --- | --- | --- | --- | --- |
| T-DM1 | 4.1 ± 0.4 | 11.2 ± 1.2 | 18.3 ± 1.1 | 21.0 ± 0.9 | 19.2 ± 0.8 | 13.4 ± 1.1 | 6.6 ± 0.3 | 3.5 ± 0.4 | 2.7 ± 0.3 | 3.4 ± 0.02 |
| TACT(T-DM1) | 3.9 ± 0.2 | 12.7 ± 0.6 | 20.0 ± 0.5 | 22.3 ± 0.4 | 18.7 ± 0.7 | 11.6 ± 0.6 | 5.4 ± 0.3 | 2.8 ± 0.2 | 2.6 ± 0.2 | 3.25 ± 0.04 |

**Supplementary Table 8. Distribution of T-DXd DAR species by intact SEC-MS post release from TACT hydrogel, compared to T-DXd reference**.

| **Conditions** | **D6 (%)** | **D8 (%)** | **avDAR** |
| --- | --- | --- | --- |
| T-DXd | 3.6 ± 0.3 | 96.4 ± 0.3 | 7.93 ± 0.01 |
| TACT(T-DXd) | 4.0 ± 0.2 | 96.0 ± 0.2 | 7.92 ± 0.01 |

**Supplementary Table 9.** Average relative peak intensities for each DAR species in the MS spectra of ADCs

|  | **Intensity (%)** | | | | | | | | |  |
| --- | --- | --- | --- | --- | --- | --- | --- | --- | --- | --- |
| **ADC** | D0 | D1 | D2 | D3 | D4 | D5 | D6 | D7 | D8 | Av. DAR |
| T-MMAE | 0 | 0 | 100 | 0 | 0 | 0 | 0 | 0 | 0 | 2 |
| T-MMAE | 0 | 0 | 18.7 | 0 | 61.5 | 0 | 19.8 | 0 | 0 | 3.8 |
| T-MMAE | 0 | 0 | 0 | 0 | 0 | 0 | 0 | 0 | 100 | 8 |

**Supplementary Table 10. Korsmeyer-Peppas and bi-exponential model kinetic parameters for the T-MMAE DAR series.**

The in vitro release kinetics of MMAE from TACT hydrogel formulations were evaluated using a panel of mathematical models, with model selection based on the Akaike Information Criterion (AIC), which penalizes model complexity relative to goodness-of-fit to favor parsimonious descriptions, complemented by root mean square error (RMSE) as a measure of absolute prediction accuracy.

| **Biexponential Model (best fit, first-order burst + sustained phases)** | | | | | | | | | | | | | | |
| --- | --- | --- | --- | --- | --- | --- | --- | --- | --- | --- | --- | --- | --- | --- |
| **Condition (DAR)** | **A_1_ (%)** | **K_1_ (h^-1^)** | | **T_1/2_,_1_ (h)** | | **A_2_ (%)** | | **K_2_ (h^-1^)** | **T_1/2_,_2_ (h)** | | **R^2^** | | **AIC** | **RMSE (%)** |
| TACT(trastuzumab), DAR 0 | 2.9 | 3.548 | | 0.20 | | 200 | | 0.0140 | 49.61 | | 0.994 | | -6.29 | 0.45 |
| TACT(T-MMAE),  DAR 2 | 1.6 | 78.205 | | 0.01 | | 63.0 | | 0.0381 | 18.19 | | 0.999 | | -24.40 | 0.17 |
| TACT(T-MMAE),  DAR 3.5 | 3.4 | 4.601 | | 0.15 | | 200 | | 0.0080 | 86.90 | | 0.994 | | -15.94 | 0.26 |
| TACT(T-MMAE),  DAR 8 | 5.5 | 1.293 | | 0.54 | | 200 | | 0.0044 | 157.96 | | 0.979 | | -7.00 | 0.43 |
| **Korsmeyer-Peppas Model** | | | | | | | | | | | | | | |
| **Condition (DAR)** | **k (% h^-n^)** | | **n** | | **R^2^** | | **AIC** | | | **RMSE (%)** | |  | | |
| TACT(trastuzumab), DAR 0 | 5.487 | | 0.681 | | 0.988 | | -4.25 | | | 0.63 | |  |  |  |
| TACT(T-MMAE),  DAR 2 | 4.079 | | 0.696 | | 0.994 | | -15.12 | | | 0.35 | |  |  |  |
| TACT(T-MMAE),  DAR 3.5 | 5.006 | | 0.502 | | 0.981 | | -10.00 | | | 0.46 | |  |  |  |
| TACT(T-MMAE),  DAR 8 | 4.594 | | 0.479 | | 0.968 | | -7.17 | | | 0.54 | |  |  |  |

**Supplementary Table 11. Mouse NCA pharmacokinetic parameters (T-DM1 and T-DXd).**

| PK parameters (NCA) | T-DM1 | | T-DXd | |
| --- | --- | --- | --- | --- |
|  | IV T-DM1 | SC TACT(T-DM1) | IV T-DXd | SC TACT(T-DXd) |
| C_max_ (mg L^-1^) | 350 | 254 | 200 | 74 |
| C_last_ (mg L^-1^) | 68.36 | 34.29 | 17.67 | 0.47 |
| T_max_ (h) | 0 | 96 | 0 | 168 |
| T_last_ (h) | 1008 | 1008 | 1008 | 1008 |
| AUC_0-tlast_ (mg h L^-1^) | 130219 | 96799 | 51983 | 21354 |
| AUC_0-inf_ (mg h L^-1^) | 185819 | 113264 | 60233 | 21422 |
| AUC_% extrap_ (%) | 29.9 | 14.5 | 13.7 | 0.32 |
| MRT_0-inf_ (h) | 820 | 516 | 488 | 223 |
| MRT_0-tlast_ (h) | 392 | 351 | 331 | 220 |
| 𝜆_z_ (h^-1^) | 0.00123 | 0.00208 | 0.00214 | 0.00690 |
| T_half_ (h) | 564 | 333 | 324 | 100 |
| T_half,eff_ (h) | 568 | 358 | 338 | 154 |
| CL (L h^-1^) | 1.88E-06 | 3.09E-06 | 3.32E-06 | 9.34E-06 |
| Vz (L) | 1.53E-03 | 1.48 E-03 | 1.55E-03 | 1.35E-03 |
| Vss (L) | 1.54E-03 | ND | 1.62E-03 | ND |
| F(%) | ND | 61% | ND | 36% |
| R^2^ | 0.754 | 0.841 | 0.941 | 0.909 |
| R^2^_ajusted_ | 0.672 | 0.788 | 0.926 | 0.879 |

ND: Not determined

**Supplementary Table 12. NHP NCA pharmacokinetic parameters.**

| PK parameters (NCA) | IV T-DM1 | | SC TACT(T-DM1) | |
| --- | --- | --- | --- | --- |
|  | Mean | SEM | Mean | SEM |
| C_max_ (mg L^-1^) | 232.51 | 19.40 | 82.94 | 13.55 |
| C_last_ (mg L^-1^) | 2.43 | 0.68 | 1.05 | 0.25 |
| T_max_ (h) | 0 | 0 | 64 | 8 |
| T_last_ (h) | 600 | 0 | 600 | 0 |
| AUC_0-tlast_ (mg h L^-1^) | 15597 | 1294 | 12092 | 1141 |
| AUC_0-inf_ (mg h L^-1^) | 16022 | 1431 | 12244 | 1183 |
| AUC_% extrap_ (%) | 2.54 | 0.68 | 1.19 | 0.28 |
| MRT_0-inf_ (h) | 151 | 12 | 143 | 4 |
| MRT_0-tlast_ (h) | 135 | 9 | 136 | 5 |
| 𝜆_z_ (h^-1^) | 5.99E-03 | 3.77E-04 | 7.24E-03 | 5.92E-04 |
| T_half_ (h) | 117 | 8 | 97 | 7 |
| T_half,eff_ (h) | 105 | 9 | 99 | 2 |
| CL (L h^-1^) | 2.14E-03 | 2.67E-04 | 2.71E-03 | 2.86E-04 |
| Vz (L) | 0.361 | 0.095 | 0.373 | 0.011 |
| Vss (L) | 0.320 | 0.055 | ND | ND |
| F(%) | ND | ND | 78.1 | 19.1 |
| R^2^ | 0.985 | 0.010 | 0.990 | 0.002 |
| R^2^_ajusted_ | 0.982 | 0.013 | 0.988 | 0.003 |

ND: Not determined

### Section 5. Supplementary References

1.	Danielsen, S. P. O. *et al.* Molecular Characterization of Polymer Networks. *Chem. Rev.* **121**, 5042–5092 (2021).

2.	Hotton, C. *et al.* Tuning Structure and Rheological Properties of Polyelectrolyte-Based Hydrogels through Counterion-Specific Effects. *Macromolecules* **56**, 923–933 (2023).

3.	Kolodych, S. *et al.* Development and evaluation of β-galactosidase-sensitive antibody-drug conjugates. *Eur. J. Med. Chem.* **142**, 376–382 (2017).
